# The transcriptomic landscape of monosomy X (45,X) during early human fetal and placental development

**DOI:** 10.1101/2024.03.01.582942

**Authors:** Jenifer P. Suntharalingham, Ignacio del Valle, Federica Buonocore, Sinead M. McGlacken-Byrne, Tony Brooks, Olumide K. Ogunbiyi, Danielle Liptrot, Nathan Dunton, Gaganjit K Madhan, Kate Metcalfe, Lydia Nel, Abigail R. Marshall, Miho Ishida, Neil J. Sebire, Gudrun E. Moore, Berta Crespo, Nita Solanky, Gerard S. Conway, John C. Achermann

## Abstract

Monosomy X (45,X) is associated with Turner syndrome and pregnancy loss in humans, but the underlying mechanisms remain unclear. We therefore analyzed the transcriptomic landscape of clinically relevant human fetal 45,X tissues (including pancreas, liver, kidney, skin, placenta) with matched 46,XX and 46,XY control samples between 11-15 weeks post conception (n=78). Although most pseudoautosomal region 1 (PAR1) genes were lower in monosomy X tissues, we also found reduced expression of several key genes escaping X inactivation (e.g., *KDM5C* and *KDM6A*), and potentially clinically important transcripts such as genes implicated in ascending aortic aneurysm. In contrast, *higher* expression of an autosomal, long non-coding RNA (*OVCH1-AS1*) was seen in all 45,X tissues. In the placenta, lower expression of *CSF2RA* was demonstrated, likely contributing to immune dysregulation. Taken together, these findings provide novel insights into the biological consequences of a single X chromosome during early human development and potential insights in genetic mechanisms in Turner syndrome.

## INTRODUCTION

Complete or partial loss of the second X (sex) chromosome in humans occurs in approximately 1:2500 girls and women and is associated with Turner syndrome (TS)^1,2^. Around 50% of individuals with TS have a monosomy X (45,X) karyotype but other variations in karyotype are often seen (e.g., isochromosome Xq (46,X,i(Xq), ring X (mosaic 45,X/46,X,r(X)), Xp or Xq deletions, and 45,X/46,XX or 45,X/46,XY mosaicism)^1–3^. Although monosomy X is the only chromosome monosomy compatible with survival in humans, it is estimated that 2% of monosomy X fetuses survive to term and many pregnancies are lost in the first or early second trimester^4–6^.

Girls and young women with TS/monosomy X can present with many different features and at different ages. In the newborn period, the diagnosis may be suspected due to lymphedema, congenital cardiovascular anomalies (e.g., coarctation of the aorta), renal features (e.g., horseshoe kidney) or distinct physical signs (e.g., wide chest, widened neck skin)^1–3,7^. Transient hyperinsulinism and hypoglycemia have also been reported^8,9^. In childhood, early features include impaired growth and recurrent otitis media, whereas absent puberty and primary ovarian insufficiency (POI) present later in teenage years^7,10,11^. Higher risks of long-term co-morbidities in adulthood are described, such as diabetes mellitus, weight gain, hypertension, raised liver enzymes, hearing impairment, hypothyroidism, autoimmunity, acquired cardiovascular disease, and skin nevi^1–3,11–15^. Women with TS have an overall increased mortality partly accounted for by aortic root dilatation and dissection^12,15–17^. Thus, the clinical features associated with TS can affect many different systems and may have some origins in early fetal development^2^. Identifying underlying mechanisms that drive the clinical features of TS is important as they may help to develop personalized medicine strategies and improve care for girls and women in the long term^1,2,18^.

Key genetic mechanisms hypothesized to drive TS phenotypes are usually related to the complete or partial loss of the second sex chromosome^1,2,19,20^. Haploinsufficiency of genes in the pseudoautosomal (PAR) regions (PAR1, PAR2), which have Y chromosome homologs, have been associated with TS phenotypes such as short stature and cubitus valgus (e.g., *SHOX*)^2,21,22^. Monosomy X may also be associated with reduced dosage of genes that normally escape X-inactivation^2,19,20,23,24^. X inactivation is a process whereby one X chromosome is transcriptionally silenced in female mammalian cells to equalize dosage of gene products from the X chromosome between 46,XX and 46,XY individuals^6,23,25^. This process is mediated largely by the non-coding RNA transcripts, *XIST*/*TSIX*^23,25^. It is well established that some genes on the X chromosome escape X inactivation; these genes are normally biallelically expressed from both X chromosomes in 46,XX females, but this may not occur in those with a 45,X karyotype^19,20,23,26,27^. Moreover, a core subset of X chromosome genes that might drive somatic sex differences has recently been proposed^25^.

In addition to direct effects of reduced PAR gene dosage, other mechanisms linked to the pathogenesis of TS include disruption of X chromosome genes that have a knock-on (or “ripple”) effect on other parts of the X chromosome itself^2,19,20^, including important non-coding RNAs such as JPX (a key regulator of the X inactivation gene, *XIST*)^19,24,28^, or X chromosome genes that influence autosomal genes with regulatory functions such as ubiquitination, chromatin modification, translation, splicing, DNA methylation and circular RNA generation^2,19,20,24,26–30^. Unravelling these complex interactions requires whole genome transcriptomic analysis at a suitable scale and in relevant tissues.

To date, most transcriptomic studies investigating the pathogenic basis of TS have analyzed blood leukocytes/peripheral blood mononuclear cells (PBMCs)^20,24,27^. Direct sampling of tissues strongly associated with phenotypic features such as diabetes, hypertension and obesity is much more challenging. Recently, transcriptomic analysis of adult fat and muscle biopsies has been reported from individuals with TS and related sex chromosome aneuploidies such as 47,XXY (Klinefelter syndrome)^19,30^. This approach is starting to provide insight into the effects of sex chromosomes in different tissues, and to identify core haploinsufficient X chromosome genes associated with a 45,X karyotype^19^. However, the biochemical and transcriptomic profile of adult tissues may also be influenced by confounding factors such as medication, inflammation, diet or the complex interplay between different systems (e.g., fat, muscle and pancreas in insulin sensitivity), so assessing the “pure” monosomy X transcriptome is difficult.

Given the fact that many clinical features associated with monosomy X are present in early postnatal life, and that the origins of many long-term adult conditions may be in part established during embryonic or fetal development^31,32^, we undertook detailed transcriptomic analysis of monosomy X fetal samples from several key tissues of interest between 11-15 weeks post conception (wpc). Our aim was to better understand the transcriptomic events associated with monosomy X in early human development, and to obtain a unique perspective on human X chromosome biology and possible disease mechanisms in Turner syndrome.

## RESULTS

### Global transcriptomic differences and key sex chromosome genes

In order to identify global transcriptomic differences associated with a single X chromosome during development, we performed bulk RNA sequencing (bulk RNA-seq) using human monosomy X fetal samples (n=20) between 11-15 wpc and compared these to tissue- and age-matched 46,XX (n=20) and 46,XY controls (n=20) (Fig. 1a, Supplementary Fig. 1 and Supplementary Data 1). Pancreas, liver, kidney, skin, and a mixed sample group (comprised of brain, heart, lung and spleen) were chosen, as these are biologically relevant to the clinical features associated with TS in childhood or in later life. All 60 tissue samples underwent single nucleotide polymorphism (SNP) array analysis on simultaneously extracted DNA. This approach confirmed the expected karyotype in all cases, except for one 45,X fetus where low-level mosaicism for a 46,XY cell line was seen (Supplementary Fig. 2).

**Figure 1.**
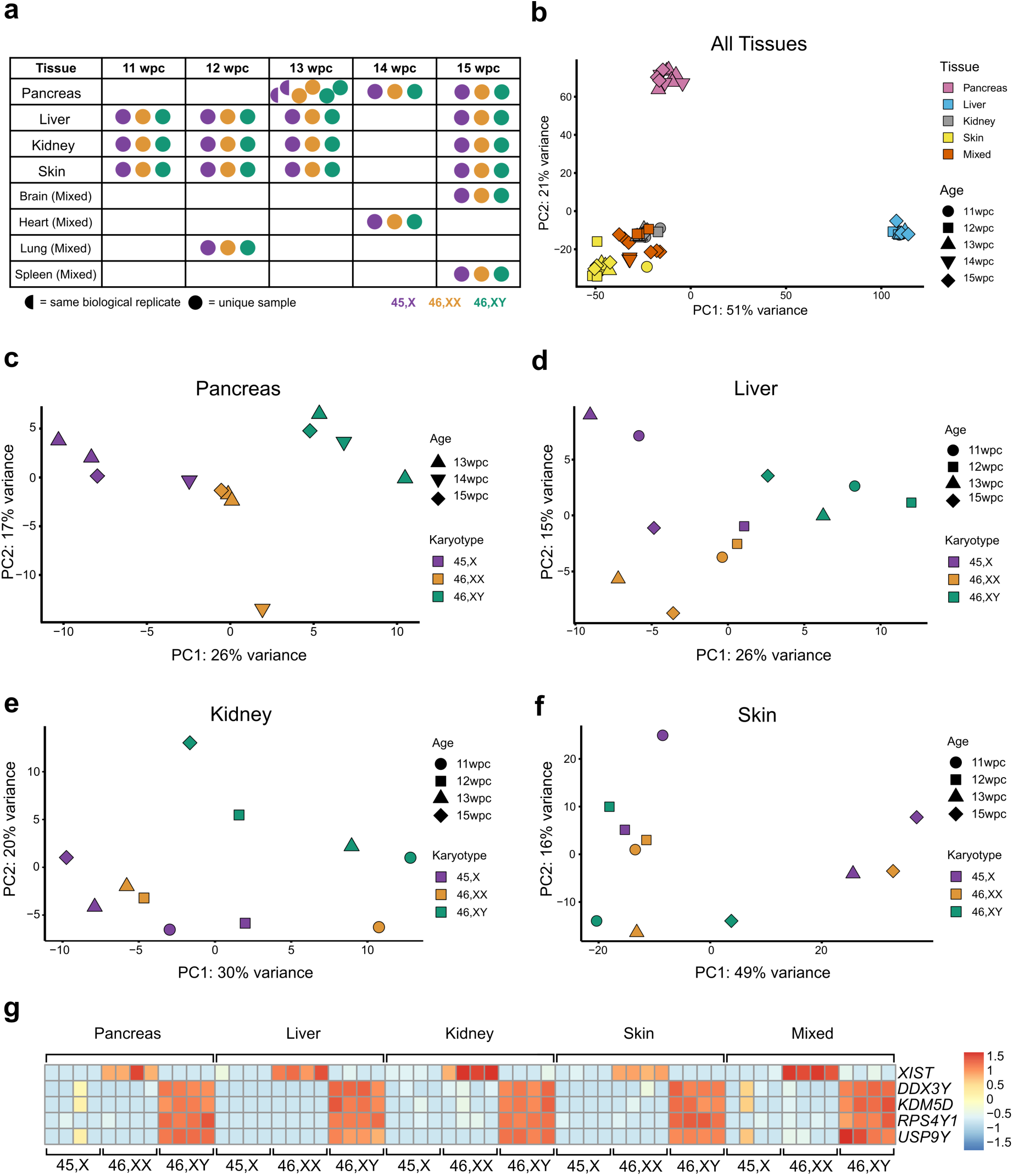
Experimental design and principal component analysis of tissues. **a** Overview of age stages and tissues used for the main study. Karyotypes are indicated by the key. One 13 week post conception (wpc) pancreas was bisected longitudinally and both parts processed independently. **b** Principal component analysis (PCA) of the whole multi-tissue dataset (n=60). PC, principal component. **c** PCA of pancreas samples (n=12). **d** PCA of liver samples (n=12). **e** PCA of kidney samples (n=11; one outlier removed due to low level (<10%) adrenal contamination). **f** PCA of skin samples (n=11; one outlier removed due to low level (<10%) muscle contamination). Age and karyotypes of tissues are shown in the key. **g** Heat map of key sex chromosome genes for each set of tissue samples; *XIST* for the X chromosome and *DDX3Y*, *KDM5D*, *RPS4Y1* and *USP9Y* for the Y chromosome. Expression intensity is shown in the key.

Using principal component analysis (PCA) for the entire dataset (n=60), samples clustered together based primarily on tissue of origin rather than karyotype, as expected (Fig. 1b, Supplementary Fig. 3 a, b). In contrast, PCA of individual tissues showed an influence of karyotype on clustering for pancreas (Fig. 1c) and liver (Fig. 1d), some effect of karyotype in kidney, and limited effect in skin (Fig.1 e, f and Supplementary Fig. 3 c, d).

Heatmap normalized expression of several important sex chromosome genes is shown in Fig. 1g. *XIST*, the principle X chromosome regulator of X inactivation, was strongly expressed in all 46,XX tissues, consistent with the presence of two chromosomes. *XIST* was not expressed in 46,XY samples nor in 45,X samples, both of which have a single X chromosome and do not undergo X inactivation. Y chromosome genes (e.g., *DDX3Y*, *KDM5D*, *RPS4Y1*, *USP9Y*) were strongly expressed in 46,XY samples and not in 46,XX samples nor 45,X samples, except at low level in the two mosaic samples (pancreas, heart/mixed) outlined above (Fig.1g).

In order to identify genes with consistently lower or higher expression in monosomy X tissues compared to controls, differential gene expression (DGE) analysis was undertaken. Volcano plots comparing 45,X samples with either 46,XX or 46,XY matched control samples for each tissue are shown in Fig. 2 (Supplementary Data 2-25). Genes with higher expression in 45,X samples have a positive log_2_ fold change (log_2_FC), whereas genes with a higher expression in either 46,XX or 46,XY samples have a negative log_2_FC. As expected, the X inactivation genes *XIST* and *TSIX* showed higher expression in 46,XX samples compared to 45,X samples, whereas Y chromosome genes were differentially expressed in 46,XY samples compared to 45,X (Fig. 2).

**Figure 2.**
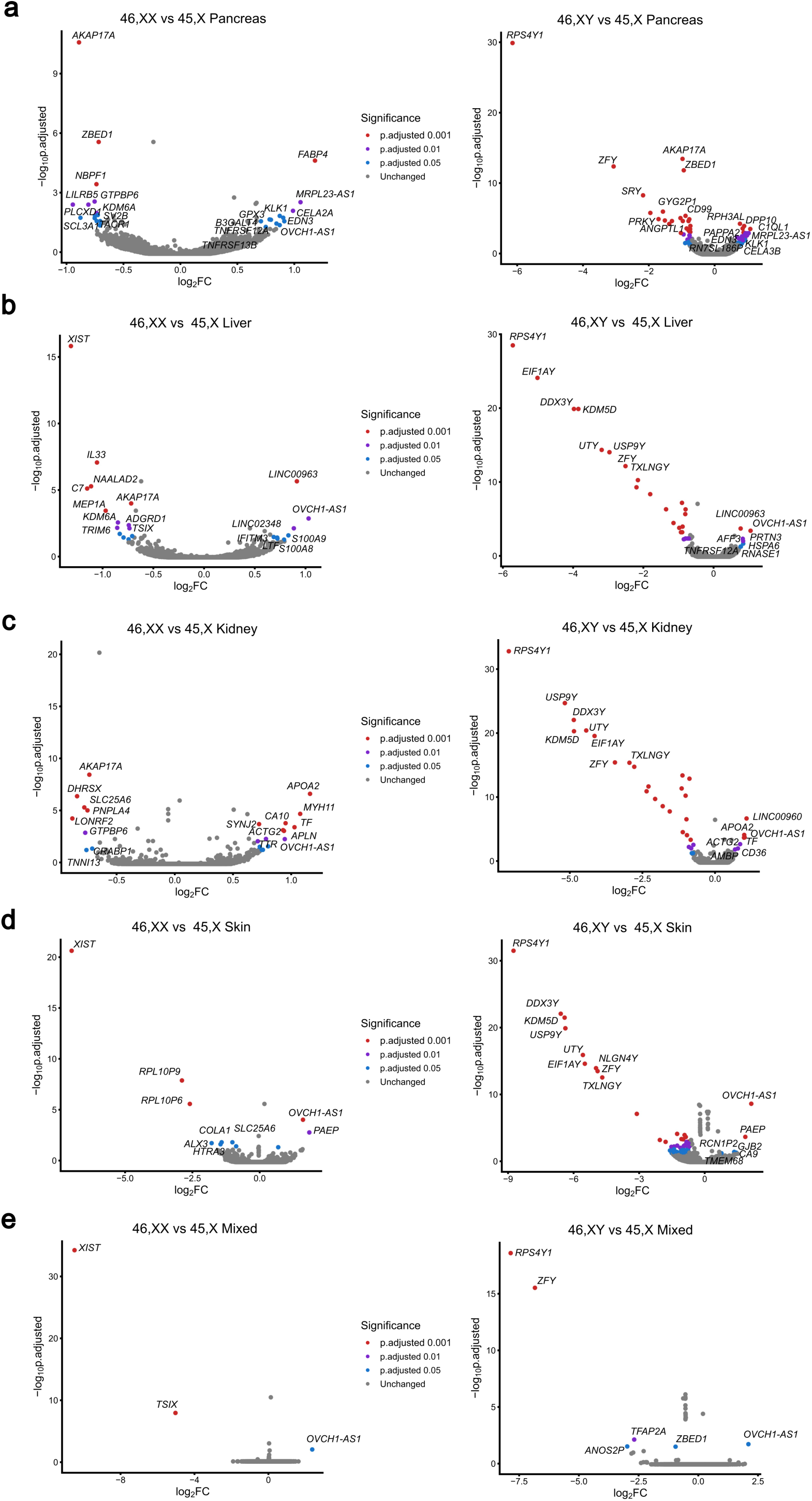
Volcano plots showing differential expression of genes between the 45,X and either 46,XX or 46,XY matched control samples. Data are shown for: **a** Pancreas; **b** Liver; **c** Kidney; **d** Skin; **e** Mixed group (brain, heart, lung, spleen). Four samples were included in each group (see experimental design Figure 1a). Comparison of 45,X *versus* 46,XX is shown in the left-hand panel and comparison of 45,X *versus* 46,XY is shown on the right-hand panel. The top ten most differentially expressed genes in each dataset are labeled, based on adjusted p-value and where log_2_ fold change (FC) is greater than +/-0.7. Genes with higher expression in 45,X tissues have a positive log_2_FC and those with higher expression in control samples have a negative log_2_FC. The significance level of highlighted points is shown in the key.

### Genes with lower expression in monosomy X tissues

Our initial analysis focused on genes that were lower in both 45,X *versus* 46,XX and 45,X *versus* 46,XY datasets, and consistent across each of the five different tissues studied (n=10 datasets) (log_2_FC<-0.5, adjusted p-value (p-adj) <0.05; Supplementary Fig. 4). This relatively low threshold for log_2_FC was chosen to identify subtle but potentially meaningful differences in gene dosage, especially related to haploinsufficiency effects in monosomy X. By intersecting these groups, no genes were shared among all tissues studied with this cut-off, although four genes (*SLC25A6*, *AKAP17A*, *GTPBP6*, *ZBED1*) did have lower monosomy X expression in multiple tissues (Fig. 3 a-c and Supplementary Fig. 4). These genes showed haploinsufficiency of expression in monosomy X tissues in violin plots (Fig. 3b and Supplementary Fig. 5). As these genes are all located in the PAR1 region of the X chromosome (Fig. 3d), investigation of the expression of all PAR genes was undertaken for 45,X *versus* 46,XX or 46,XY tissues (Fig. 3e). This analysis showed consistently lower expression of many PAR1 genes in 45,X tissues compared to controls, where log_2_FC=-1.0 represents haploinsufficiency in 45,X, and where log_2_FC=0 represents similar expression in control and 45,X samples (Fig. 3e). Several PAR1 genes did not show differential expression; these genes generally had low transcript counts, or were not detected in all tissues (e.g., *SHOX*, *CRLF2*, *P2RY8*, *ASMT*, *XG*) (Supplementary Fig. 5 and Supplementary Data 26 and 28). Of note, the three PAR2 region genes (*SPRY3*, *VAMP7*, *IL9R*) also did not show marked differential expression (Fig.3e, Supplementary Fig. 6, and Supplementary Data 27 and 28), as shown in samples from adults with monosomy X^19,20,30^.

**Figure 3.**
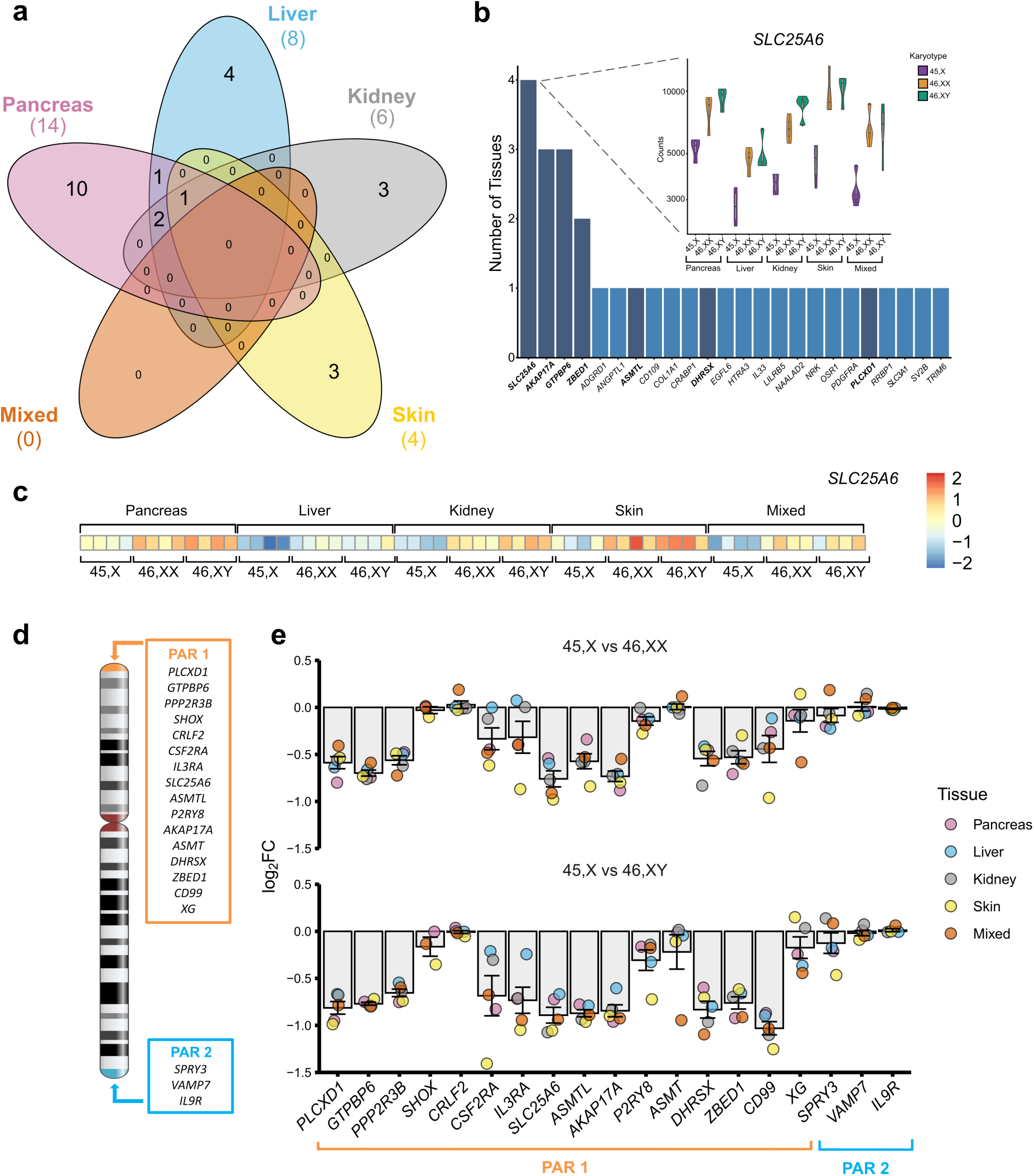
Genes with lower expression in monosomy X when compared to 46,XX or 46,XY controls. **a** Five-way Venn diagram showing overlap of genes with lower expression in monosomy X when compared to control karyotypes across all tissues. **b** Graph to show genes with lower expression in monosomy X across all tissues with inset of violin plot showing normalized count data of *SLC25A6* across all tissues (from panel a) according to karyotype (n=4 in each group). Pseudoautosomal region (PAR) genes are shown in bold and dark bars. **c** Heatmap of *SLC25A6* expression across all tissues used in study. **d** Schematic of the X chromosome and showing PAR1 and PAR2 genes. **e** Differential expression of genes located on the PAR regions for 45,X compared to 46,XX or 46,XY samples. Individual mean data points for each tissue group are shown, as indicated in the legend. The bars represent the mean of the different tissue groups with standard error of the mean shown. Log_2_ fold change (FC)=-1.0 represents half the expression in 45,X samples (i.e., haploinsufficiency), whereas log_2_FC=0 represents similar expression in 45,X samples and 46,XX and 46,XY controls (n=4 for each karyotype in each tissue group).

In addition to PAR genes, recent reports have identified core sets of genes that likely escape X inactivation or influence phenotype in monosomy X adult tissues^19^, as well as ten X chromosome genes that are implicated as key mediators of biological sex differences^25^. Analysis of these genes in monosomy X tissues during development compared to controls is shown in Fig. 4, a-c, and Supplementary Data 29-30. Notably, the histone demethylase genes, *KDM5C*/*JARID1C*/*SMCX* and *KDM6A*/*UTX* had lower expression in 45,X compared to 46,XX control tissues, likely due to X inactivation escape. Although similar expression of these two genes was seen between 45,X and 46,XY tissues, relative haploinsufficiency of demethylase activity is still likely to occur in monosomy X compared to 46,XY tissues as 46,XY tissues have compensation by their Y chromosome homologues, *KDM5D*/*JARID1D*/*SMCY* and *UTY*/*KDM6C*, respectively. Thus, histone methylation status may be altered in 45,X and could influence developmental processes.

**Figure 4.**
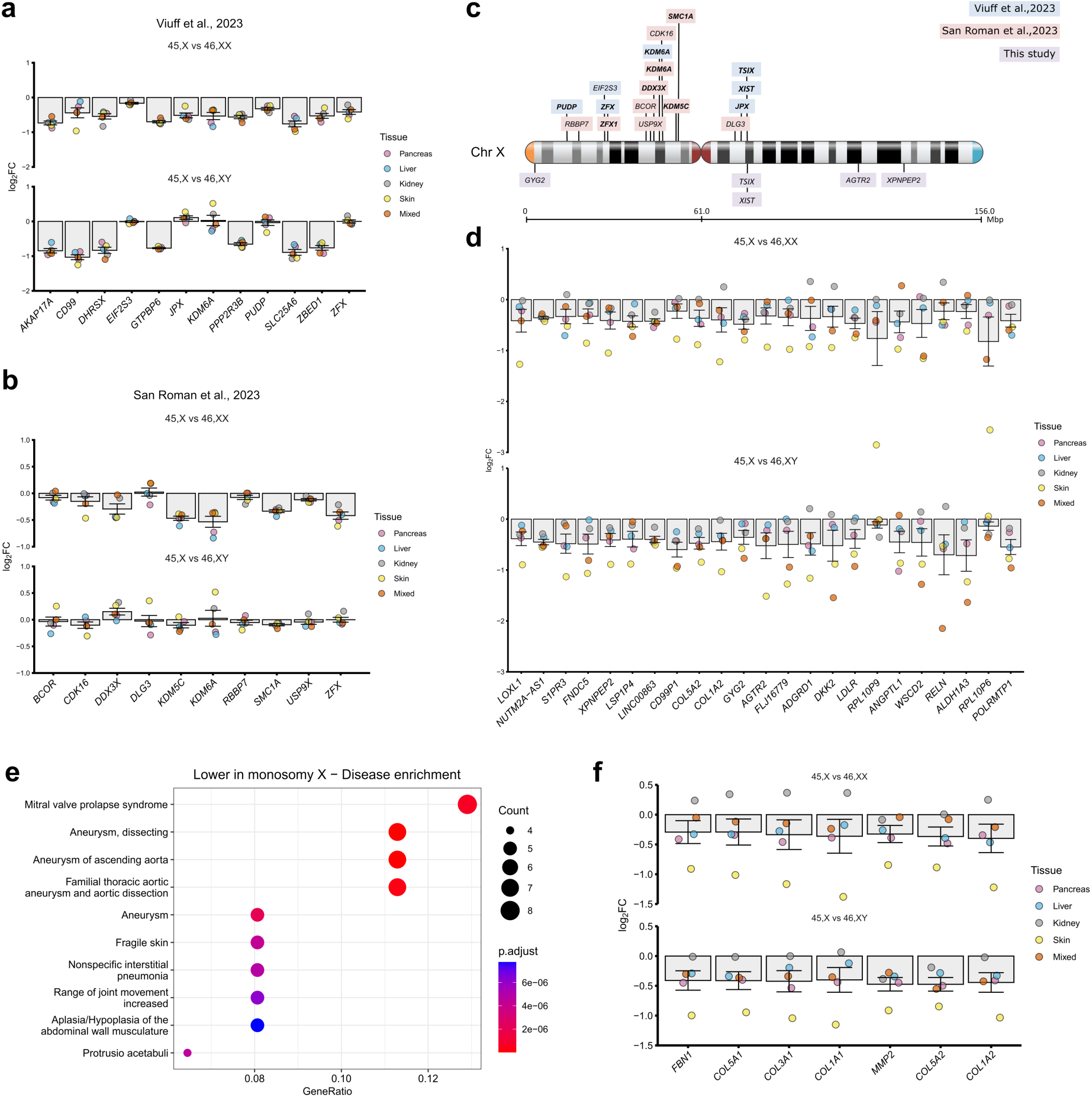
Additional genes that have lower expression in monosomy X samples and related pathways. **a** Differential expression in our datasets of selected core X chromosome genes linked to monosomy X, including those linked to X chromosome dosage (from Viuff et al, 2023)^19^. Individual mean data points for each tissue group are shown, as indicated in the legend. The bars represent the mean of these different tissue groups with standard error of the mean shown. Log_2_ fold change (FC)=-1.0 represents half the expression in 45,X samples (i.e., haploinsufficiency), whereas log_2_FC=0 represents similar expression in 45,X samples and 46,XX and 46,XY controls (n=4 for each karyotype in each tissue group). Note *XIST* and *TSIX* show strong differential expression in all datasets but are omitted from the graphic. **B** Differential expression in our datasets of selected X chromosome genes linked to sex differences (from San Roman et al, 2023)^25^. Data are presented as described above. Note the different y-axis scale compared to Fig. 4a. **c** Schematic showing the genomic location of key genes on the X chromosome. PAR genes are not shown. **d** Differential expression data of other X chromosome and autosome genes that are lower in 45,X compared to 46,XX or 46,XY samples. Data are presented as described above and ordered according to the mean of the 45,X *versus* 46,XX and 45,X *versus* 46,XY datasets. **e** Dot plot showing disease enrichment analysis for genes lower in 45,X compared to mean controls (n=74), with log_2_FC cut-off below −0.35. **f** Differential expression in our dataset of the key genes contributing to the thoracic aortic aneurysm disease enrichment pathway. Data are presented as described above.

Next, we analyzed genes that showed global lower expression in monosomy X across multiple tissues (total n=10; of which n=5 were 45,X<46,XX and n=5 were 45,X<46,XY). We hypothesized that these genes could also have important biological functions, beyond the PAR/XCI model. In addition to the PAR genes and XCI escape genes described above, several notable autosomal genes (e.g., *ALDH1A3*, *RELN*, *LDLR*, *DKK2*) and other X chromosome genes (e.g., *AGTR2*) emerged as having lower expression in 45,X tissues (mean log_2_FC below −0.4 for 45,X compared to controls shown in Fig. 4d and Supplementary Data 31). Furthermore, when a log_2_FC below −0.35 cut off was applied to identify key genes for disease enrichment pathway analysis (n=74 genes), biological functions emerged related to dissecting aneurysm, aneurysm of the ascending thoracic aorta and connective tissue disorders (Fig. 4e). These functions resulted largely from lower expression of core connective tissue genes in monosomy X (i.e., *FBN1*, *COL5A1*, *COL3A1*, *COL1A1*, *MMP2*, *COL5A2*, *COL1A2*) (Fig. 4f). The mean differences in gene dosage were subtle, and not all values were significant when adjusted for multiple comparisons, but more marked differences were seen in the skin where collagen genes are innately expressed (Fig. 4f). We hypothesize, therefore, that lower expression of these genes could confer a risk for the development and progression of aortic root dilatation and aortic dissection in women with TS, especially as loss of *FBLN* is found in Marfan Syndrome where thoracic aortic aneurysm can be a key feature^33^.

### Genes with higher expression in monosomy X tissues

Although lower expression of many X chromosome genes was expected in monosomy X tissues, especially in the PAR1 region, we also addressed whether any X chromosome or autosome genes show consistently *higher* expression in monosomy X tissues compared to matched controls.

In order to achieve this, a tissue specific analysis was first performed to identify genes that were higher in both 45,X *versus* 46,XX and 45,X *versus* 46,XY datasets (log_2_FC>0.5, p-adj ≤0.05; Supplementary Fig. 7). Next, the intersect of these genes across all five tissue groups was generated (Fig. 5 a, b).

**Figure 5.**
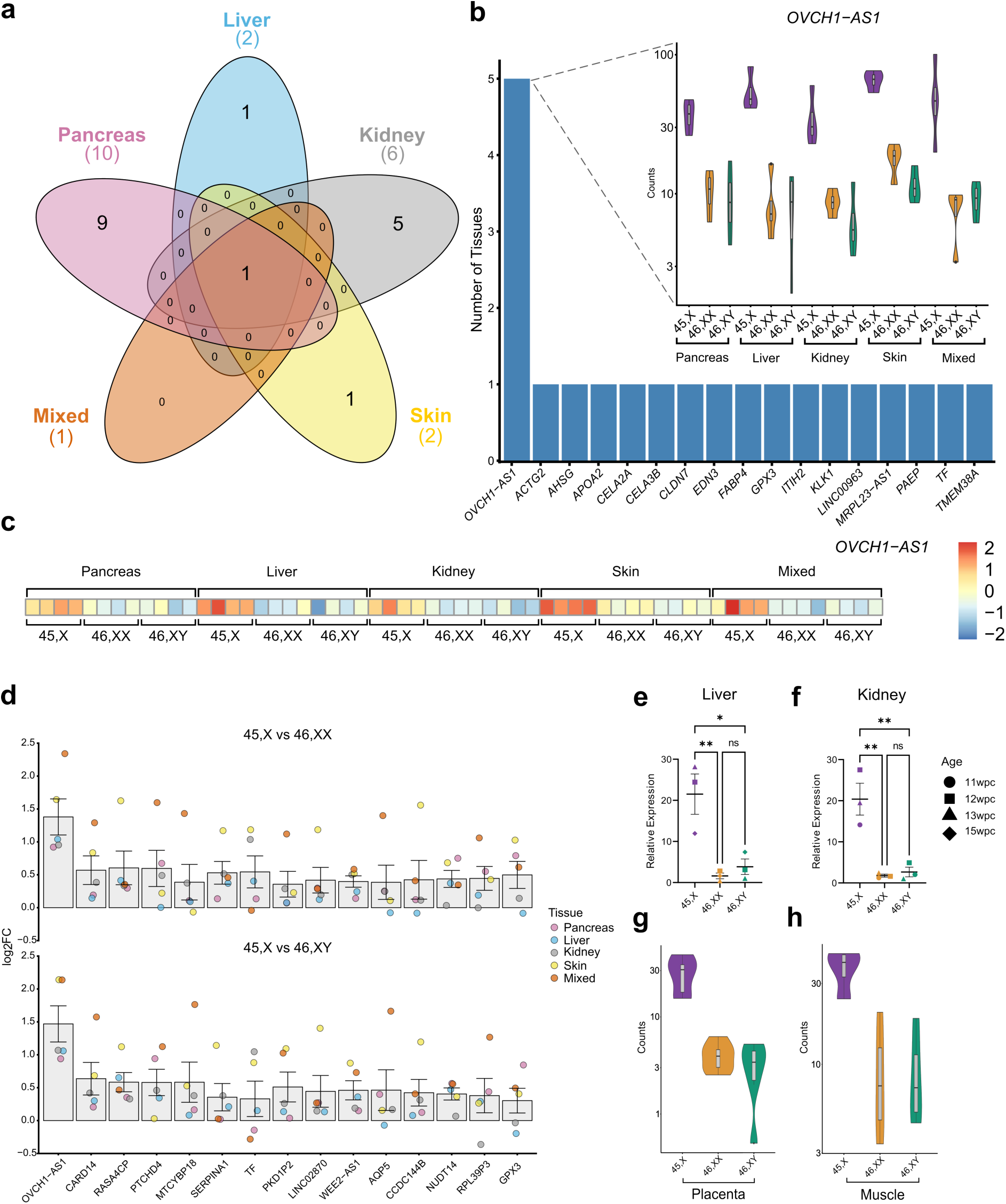
Genes with higher expression in monosomy X when compared to 46,XX or 46,XY controls. **a** Five-way Venn diagram of differentially expressed genes with higher expression in monosomy X tissues. **b** Graph to show genes with higher expression in 45,X with inset showing violin plot of normalized count data of *OVCH1-AS1* across all tissues according to karyotype (n=4 in each group). **c** Heat map of *OVCH1-AS1* expression across all tissues used in the study. **d** Differential expression data of genes that are higher in 45,X compared to 46,XX or 46,XY samples. Individual data points for each tissue are shown, as indicated in the legend. The bar represents the mean of these data points with the standard error of the mean shown. Here, log_2_ fold change (FC)=1.0 represents twice the expression in 45,X tissues, whereas log_2_FC=0 represents similar expression in 46,XX and 46,XY control and 45,X samples. (n=4 for each karyotype in each tissue group). **e** Quantitative real-time PCR (polymerase chain reaction) showing relative expression of *OVCH1-AS1* in fetal liver (n=4 samples in each group). Data are presented as scatter dot plots with each datapoint being the mean of triplicate experiments. The mean and standard error of the mean of these datapoints is also shown. **f** Quantitative real-time PCR showing relative expression of *OVCH1-AS1* in fetal kidney (n=4 samples in each group). Data are presented as scatter dot plots with each datapoint being the mean of triplicate experiments. The mean and standard error of the mean of these datapoints is also shown. **g** Violin plots of *OVCH1-AS1* expression (normalized counts) in 45,X placenta compared to 46,XX and 46,XY controls (n=6 in each group). **h** Violin plots of *OVCH1-AS1* expression (normalized counts) in 45,X muscle compared to 46,XX and 46,XY controls (n=4 in each group). p-value <0.05; **, p-value <0.01; ns, not significant.

Remarkably, only one gene was consistently identified across all tissues (Fig. 5a). This gene, OVCH1 Antisense RNA 1 (*OVCH1-AS1*) had higher expression in all monosomy X tissues, including the mixed group, when compared to 46,XX and 46,XY controls (Fig. 5 b-d; see also Fig. 2, Supplementary Fig. 7, and Supplementary Data 32 and 33). These findings were confirmed using qRT-PCR (Fig. 5 e, f), and in replication datasets of bulk RNA-Seq from placental tissue and muscle (Fig. 5 g, h).

*OVCH1-AS1* is a long non-coding RNA (lncRNA) gene located on chromosome 12 (UCSC, chromosome12:29,389,642-29,487,473, human GRCh38/hg38) (Fig.6 a, b). *OVCH1-AS1* has been identified as a potential antisense transcript of the protein coding gene, ovochymase 1 (*OVCH1*), and maps to a locus containing *FAR2* (forward strand) and *TMTC1* and *ERGIC2* (reverse strand) (Fig. 6a).

**Figure 6.**
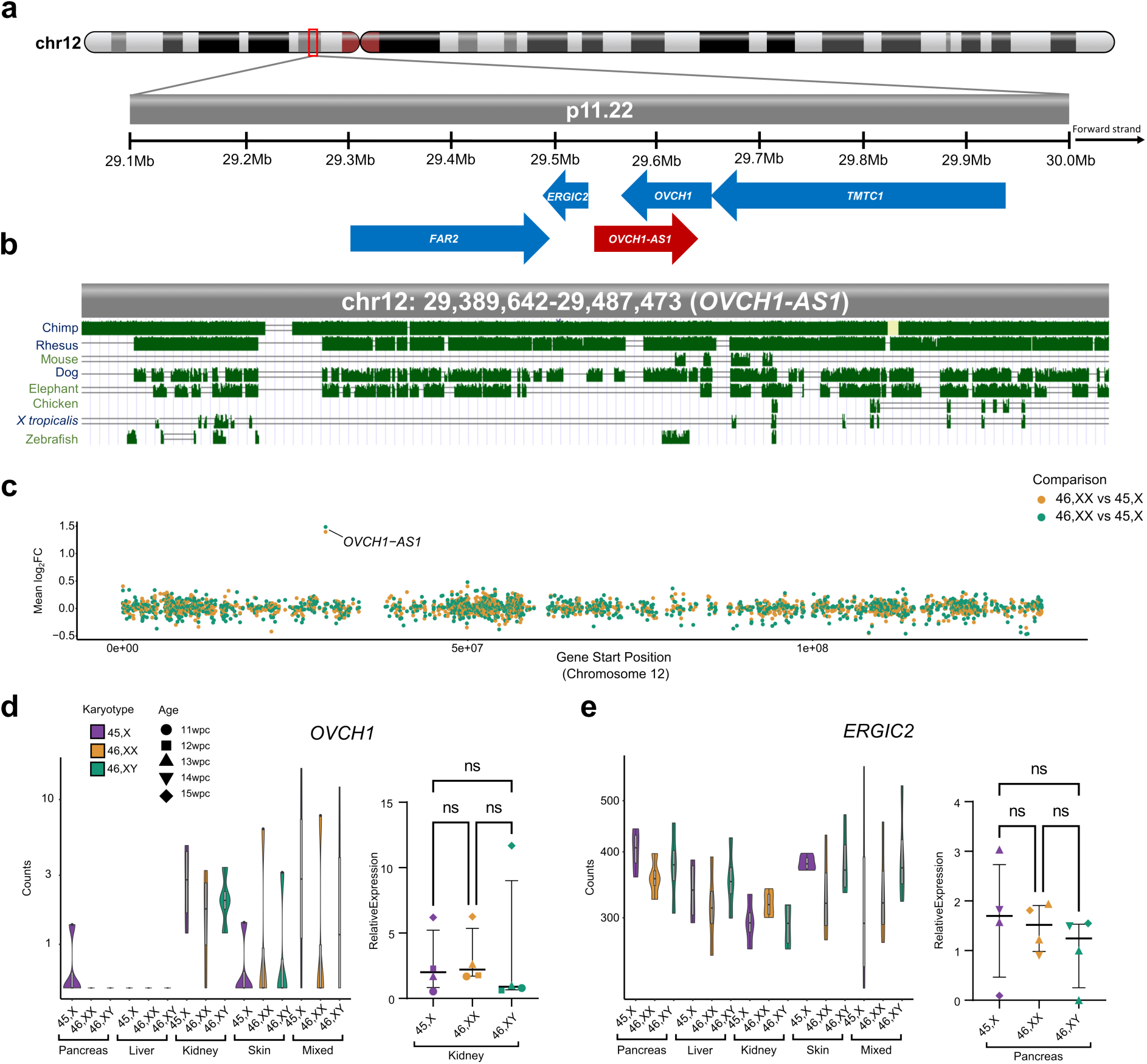
OVCH1-AS1 locus and conservation. **a** Overview of chromosome 12 (p.11.22) and genes located in the region of *OVCH1-AS1*. Arrows indicate direction of transcription. The red arrow indicates *OVCH1-AS1* (non-coding RNA gene) and blue arrows indicate protein coding genes. **b** Multiz alignments of species across the *OVCH1-AS1* locus. A single line indicates no transcript in the aligned species. Double lines show aligning species having one or more unalignable transcripts in the gap region. **c** Relative expression of all chromosome 12 genes in 45,X tissues compared to 46,XX (orange) and 46,XY (green). The higher expression of *OVCH1-AS1* in monosomy X tissue is indicated as log2 fold change (FC) in relation to control tissues. **d** Violin plot of *OVCH1* in the different tissues (normalized counts from bulk RNA-sequencing) (left panel) (n=4 samples in each group). Quantitative real-time PCR (polymerase chain reaction) showing relative expression of *OVCH1* in fetal kidney (right panel) (n=4 samples each group). Data are presented as scatter dot plots with mean and standard error of the mean also shown. **e** Violin plot of *ERGIC2* in different tissues (bulk RNA-seq) (left panel) (n=4 samples in each group). Quantitative real-time PCR showing relative expression of *ERGIC2* in fetal pancreas (right panel) (n=4 samples each group). ns, not significant.

Relatively little is known about *OVCH1-AS1*, but new studies are emerging^34^. In the GTEx database of adult tissue gene expression, *OVCH1-AS1* generally has low level expression in adult tissues, as is often the case with lncRNAs (Supplementary Fig. 8). Conservancy analysis in vertebrates using Multiz alignments (UCSC genome browser) showed that *OVCH1-AS1* is not highly conserved beyond higher primates; for example, it is not identified in mouse (Fig. 6b). Although a potential open reading frame exists between nucleotides 231-825 (203 codons) using the Coding-Potential Assessment Tool (CPAT) (RNAcentral, transcript variant 1 (URS000075D789_9606; CPAT coding probability 0.824, cut-off of 0.364), most other algorithms suggest *OVCH1-AS1* (RefSeq ID: NR_073172) is a lncRNA rather than a protein coding gene (Supplementary Table 1).

As lncRNA and anti-sense RNA transcripts can potentially modulate gene transcription, we analyzed the relative expression of all chromosome 12 genes in our dataset (45,X *versus* 46,XX and 45,X *versus* 46,XY, independently). We did not identify consistently altered gene expression in this region (Fig. 6 a-c). Next, more granular analysis of expression of genes in the *OVCH1-AS1* locus was undertaken (*OVCH1*, *ERCIG2*, *FAR2*, *TMTC1*) using bulk RNA-seq data for each tissue as well as qRT-PCR. No significant differences in the relative expression of these genes were observed between monosomy X samples and the 46,XX and 46,XY controls in these tissues studied (Fig. 6 d, e, and Supplementary Fig. 9 a, b, and Supplementary Data 34). Thus, *OVCH1-AS1* is clearly a consistently upregulated autosomal long non-coding RNA associated with monosomy X in multiple tissues, but its potential biological role as a mediator of phenotypes in women with TS requires further investigation.

### Monosomy X and the developing placenta

Although monosomy X is the only chromosomal monosomy compatible with survival in humans, it has been estimated that 98% of monosomy X pregnancies are lost^4–6^. Several hypotheses have suggested that this could be due to fetal anomalies (such as lymphedema or hydrops fetalis), defects in placental development and function, or mechanisms involving the maternal-fetal interface^4,35,36^. Detailed transcriptomic analysis of human monosomy X placenta is lacking, especially during the early stages of fetal development when loss of many monosomy X pregnancies occurs.

In order to investigate this further at a biologically relevant time point (late first trimester/early second trimester), samples were taken from monosomy X placentas (n=6) between 11-15 wpc, together with matched 46,XX (n=6) and 45,XY (n=6) control placental tissues (Fig. 7a, Supplementary Fig. 1a, and Supplementary Fig.10a).

**Figure 7.**
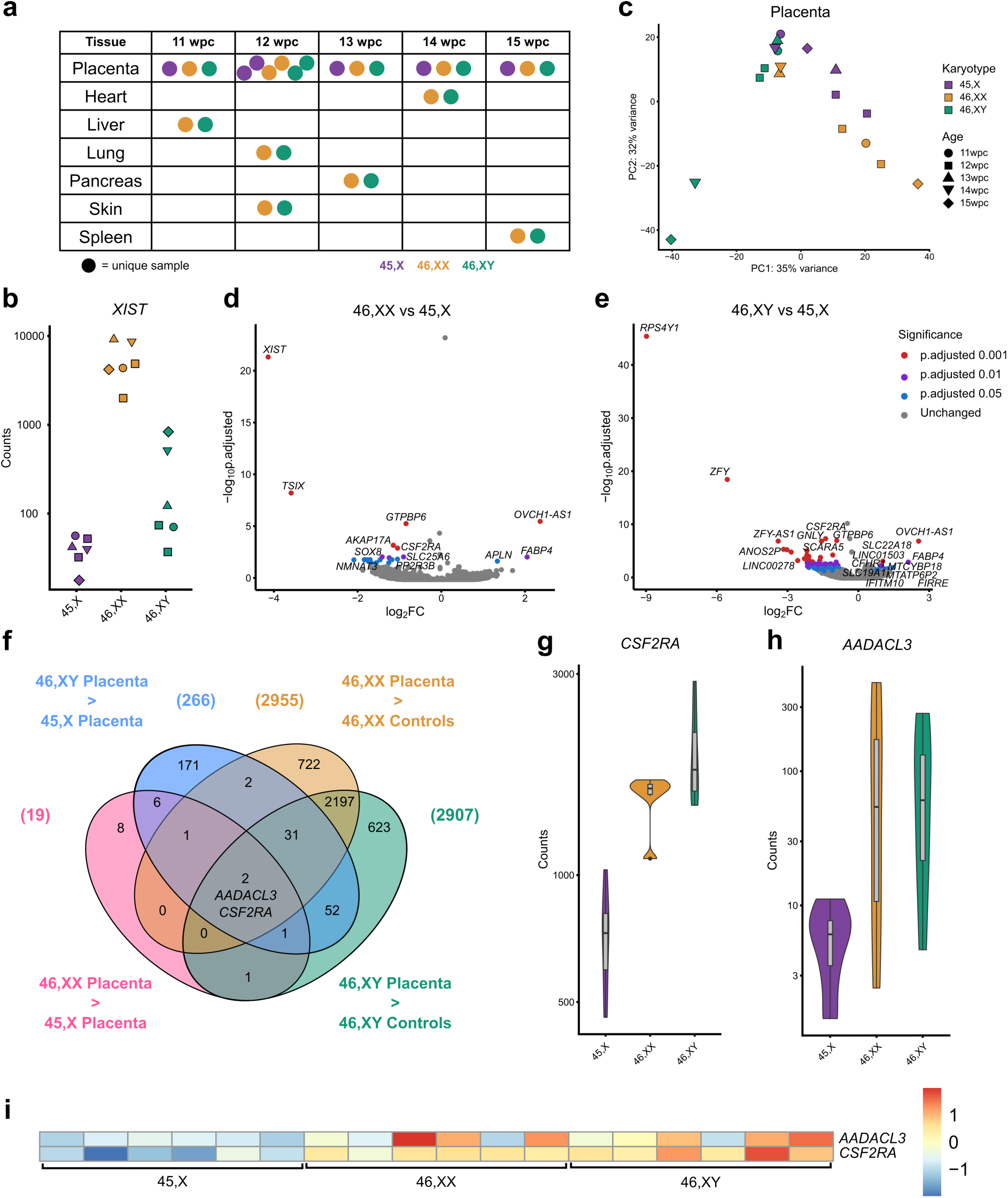
Differential expression of genes in the 45,X placenta compared to 46,XX or 46,XY controls. **a** Overview of placental age stages used for the study. Karyotypes are indicated by the key. wpc, weeks post conception. **b** Expression of the X inactivation regulator *XIST*, bulk RNA-sequencing (RNA-seq) counts in the 45,X placenta group (n=6) compared to 46,XX placenta (n=6) and 46,XY placenta (n=6). **c** Principal component analysis (PCA) of 45,X, 46,XX and 46,XY placental samples used in the study. PC, principal component. **d** Volcano plot showing differential expression of genes between the 45,X and 46,XX matched control samples (n=4 in each group). The top ten most differentially expressed genes in each dataset are labeled, based on adjusted p-value (p-adj) and where log_2_ fold change (FC) is greater than +/-0.7. Genes with higher expression in 45,X samples have a positive log_2_FC and those with higher expression in control tissues have a negative log_2_FC. The significance level of highlighted points is shown in the key. **e** Volcano plot showing differential expression of genes between the 45,X and 46,XY matched control samples. **f** Venn diagram with the common intersection showing placenta-specific genes that are consistently lower in 45,X. Data were generated for each group (n=6 versus n=6) with a differential expression cut-off of log_2_FC>0.5 and p-adj <0.05. **g** Violin plot of *CSF2RA* expression (normalized counts) in the placenta (bulk RNA-seq) (n=6 each group). **h** Violin plot of *AADACL3* expression (normalized counts) in the placenta (bulk RNA-seq) (n=6 each group). **i** Heat map of *AADACL3* and *CSF2RA* expression across all placental samples (n=6 samples in each group).

As placental mosaicism for a fetal 46,XX cell line has been proposed as a mechanism that can “rescue” or modify placental disruption^36^, we initially undertook SNP array analysis using DNA derived from two independent areas of each placenta in the 45,X group (n=6, total n=12 areas), as well as 46,XX and 46,XY controls. As expected, a low level 46,XX component was identified in all samples (45,X; 46,XY), consistent with the presence of limited maternal decidual tissue (Supplementary Fig. 11). Strong enrichment of a 46,XX line was not seen in 45,X samples compared to 46,XY controls. These findings were validated by investigating the expression of *XIST* in bulk RNA-seq data from 45,X samples, where similar transcript levels to 46,XY controls were seen (Fig. 7b and Supplementary Fig. 10b). Taken together, these data suggest that placenta mosaicism for a 46,XX cell line is not common in 45,X placenta.

We then analyzed bulk RNA-seq data to identify genes that were lower in 45,X placenta (n=6) compared to controls, which could help to elucidate potential mechanisms of pregnancy loss associated with monosomy.

Global transcriptomic analysis using PCA showed that placental samples clustered together compared to control tissues as expected (Supplementary Fig. 12), and that within the placenta samples an influence of karyotype and age was seen (Fig. 7c). Volcano plots comparing 45,X with 46,XX and 46,XY placental samples again revealed the expected pattern of *XIST* and Y chromosome gene changes, with higher expression of *OVCH1-AS1* also seen in 45,X tissue (Fig. 7d, e, and Supplementary Data 35-38). To investigate placenta-specific genes that were consistently lower in 45,X placenta, a four-way analysis was undertaken using datasets derived from 46,XX or 46,XY *versus* 45,X placenta (“45,X lower” genes), as well as from 46,XX or 46,XY placenta *versus* 46,XX or 46,XY control tissues (“placenta-specific” genes) (Fig. 7f, Supplementary Data 39-40). Using this approach, the PAR1 gene *CSF2RA* (UCSC, chrX:1,268,814-1,309,935 41,122, human GRCh38/hg38) was identified as the principal placenta-specific gene that is lower in monosomy X (Fig. 7 f-i, and Supplementary Data 41). Lower expression of an autosomal gene *AADACL3* (UCSC, Chr1:12,716,110-12,728,760, human GRCh38/hg38) was also seen, although expression levels across samples were lower and more variable (Fig.7f, h, and i, and Supplementary Data 41).

*AADACL3* encodes arylacetamide deacetylase-like3, a potential membrane protein with hydrolase activity. This gene has low level expression in adult tissues and is only found in placenta, skin and breast (Human Protein Atlas Consensus RNA-seq) (Supplementary Fig. 13a). Consistent with this, analyses of available single cell RNA-sequencing (scRNA-seq) datasets^37–39^ show low level expression of *AADACL3* in the extravillous trophoblast of early placenta, and in cytotrophoblasts, stromal cells, endothelial cells and decidual cells in late third trimester and pre-term tissue (Supplementary Fig. 14c and Supplementary Fig. 15b)^37–39^. The role of *AADAC3L* and its protein in the placenta is unknown.

In contrast, *CSF2RA* shows much stronger and more specific expression in placenta, as well as in adult haematopoietic/immune regulating tissues (bone marrow, lymph node, tonsil, spleen) (Human Protein Atlas Consensus RNA-seq) (Supplementary Fig. 13b). *CSF2RA* encodes the Colony Stimulating Factor 2 Receptor Subunit Alpha (also known as Granulocyte-Macrophage Colony-Stimulating Factor (GM-CSF) Receptor Subunit Alpha or CD116). This protein forms part of a heterodimeric cytokine receptor complex that mediates the effects of colony stimulating factor 2 (CSF2)^40^. CSF2 (GM-CSF) acts through this low affinity receptor via STAT5 to stimulate the proliferation, differentiation and functional activation of hematopoietic cells^40^. In available single cell RNA sequencing (scRNA-seq) data from first trimester placenta, *CSF2RA* expression was observed in placental villous cytotrophoblast, extravillous trophoblast and syncytiotrophoblast cells, and well as in decidual macrophages (Fig. 8a and Supplementary Fig. 14d)^37^. More granular analysis of term placenta by scRNA-seq shows high expression of *CSF2RA* in many hematopoietic cell lineages (macrophages, NK-cells, activated and resting T-cells), as well as in non-proliferative interstitial cytotrophoblasts (Supplementary Fig. 15c)^38^.

**Figure 8.**
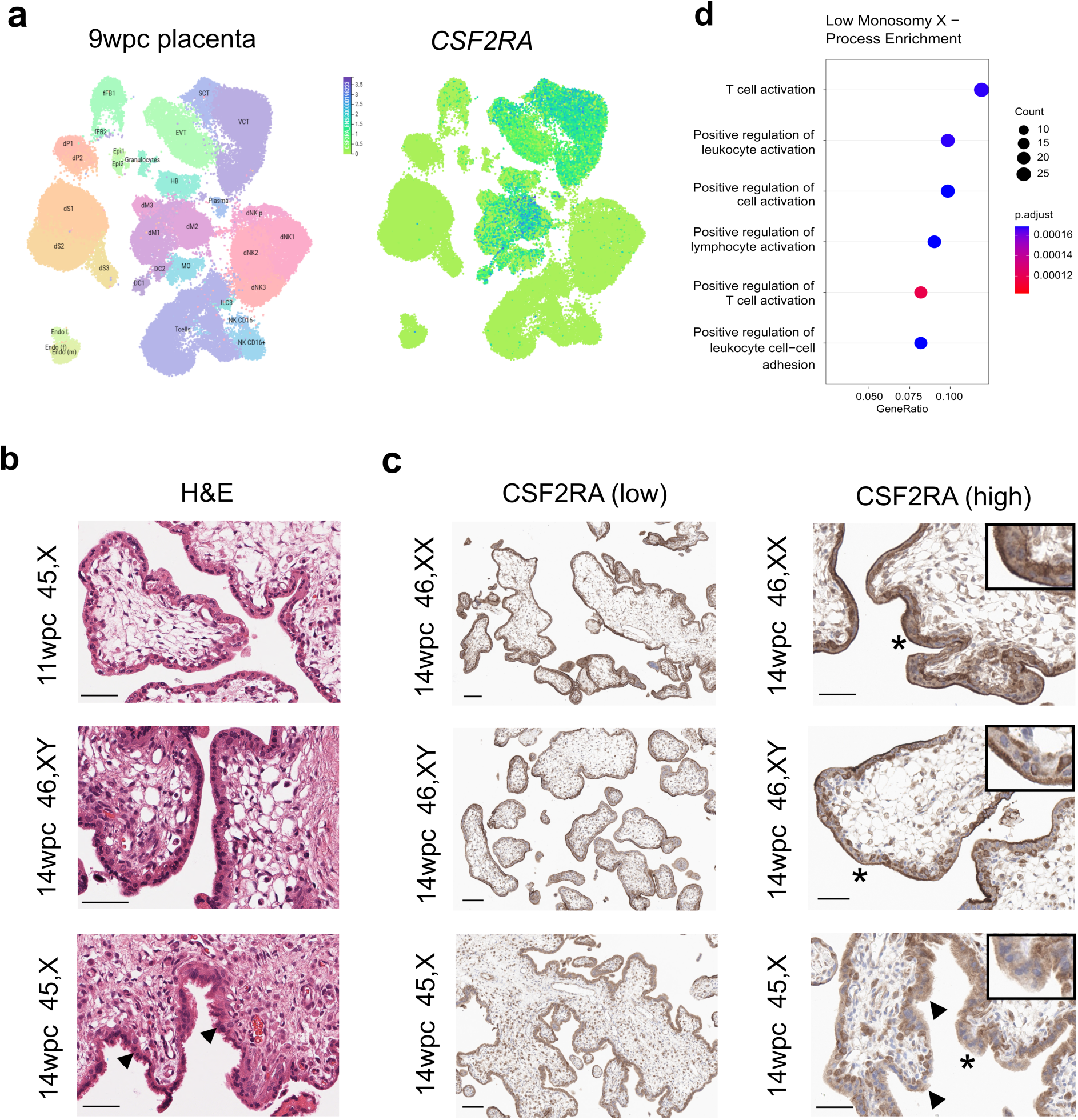
Expression of CSF2RA in the placenta and related pathways. **a** Consensus first trimester single cell data Uniform Manifold Approximation and Projection (UMAP) for clusters based on cell type (*left panel*) and feature plot for *CSF2RA* (*right panel*). These data were generated by the Vento-Tormo/Teichmann groups at the Wellcome Sanger Institute, Hinxton, UK and can be accessed using CZ CELLxGENE from https://maternal-fetal-interface.cellgeni.sanger.ac.uk/ (Vento-Tormo, R., Efremova, M., Botting, R.A. et al. Single-cell reconstruction of the early maternal–fetal interface in humans. Nature 563, 347–353 (2018); https://doi.org/10.1038/s41586-018-0698-6). This graphic is generated under a Creative Commons Attribution-BY 4.0 International License (https://creativecommons.org/licenses/by/4.0/). DC, dendritic cells; dM, decidual macrophages; dS, decidual stromal cells; Endo, endothelial cells; Epi, epithelial glandular cells; EVT, extravillous trophoblast; f, fetal; F, fibroblasts; HB, Hofbauer cells; ILC, innate lymphocyte cells; l, lymphatic; m, maternal; M3, maternal macrophages; PV, perivascular cells; p, proliferative; SCT, syncytiotrophoblast; VCT, villous cytotrophoblast. **b** H&E staining of 45,X placenta at 11 weeks post conception (wpc) (*upper panel*), 46,XY placenta at 14 wpc (*center panel*) and 45,X placenta at 14 wpc) (*lower panel*) (scale bars 50 μm). Arrowheads show regions of irregular villus border in the 14wpc 45,X sample. **c** Immunohistochemistry of CSF2RA (Colony Stimulating Factor 2 Receptor Subunit Alpha) in 46,XX, 46,XY and 45,X placenta at 14wpc at lower magnification (*left panels*) (scale bars 100 μm) and at higher magnification (*right panels*) (scale bars 50 μm). Arrowheads show regions of irregular border and diffuse CSF2RA staining in the 14wpc 45,X villus. Inset images show higher magnification of regions indicated by an asterix. **d** Biological process enrichment analysis for genes lower in 45,X placenta compared to 46,XY controls (bulk RNA-sequencing n=6 each group; 266 genes identified) with a differential expression cut-off of log_2_FC>0.5 and adjusted p-value <0.05.

To obtain more direct evidence for the role of *CSF2RA* in the monosomy X placenta, we undertook immunohistochemistry (IHC) for CSF2RA in 45,X placenta compared to 46,XX and 46,XY control samples. Between 11wpc and 15wpc the monosomy X placenta shows increasing non-specific inflammatory changes as well as irregular villus tufting (Fig. 8b). CSF2RA localized to the syncytio- and cyto-trophoblastic layers in all samples as expected, but in the 45,X placenta at 14wpc showed less defined surface staining (Fig. 8c). Furthermore, pathway enrichment analysis of genes (n=266) that are lower in monosomy X placenta compared to control (46,XY) identified processes linked to leukocyte activation and adhesion, immune response, and cytokine signaling (Fig. 8d).

Taken together, these data provide some of the first direct evidence that CSF2RA is altered in monosomy X placenta, and these findings suggest that dysregulation of immune and inflammatory processes may contribute to the placental dysfunction associated with loss of an X chromosome.

## DISCUSSION

Monosomy X is the most common chromosomal aneuploidy in humans and can be associated with a broad range of clinical features in Turner syndrome. Some of these features have a developmental origin. The exact mechanisms that give rise to phenotypes in women with TS are still poorly understood, as are the mechanisms responsible for the high incidence of pregnancy loss. We therefore undertook a detailed transcriptomic analysis of monosomy X tissues during early human development, to better understand the molecular basis of Turner syndrome, with the ultimate aim of developing personalized approaches to long-term management.

We first assessed genes with lower expression in monosomy X, to be consistent with the long-standing hypothesis that haploinsufficiency of genes in the X chromosome may drive phenotypic features. As expected, many of these genes were pseudoautosomal genes (e.g., *SLC25A6*, *AKAP17A*, *GTPBP6*, *ZBED1*, *ASTML*, *DHRSX*, *PLCXD1*) expressed from the PAR1 region. Not all PAR genes showed lower expression, mostly due to lower background transcript levels or greater variability in expression between different tissues (e.g., *SHOX*). Notably, we found PAR2 genes had no differences in expression between monosomy X and control karyotypes. This finding has also been shown in other recent studies^19,20,30^, suggesting that the dynamics of X chromosome PAR2 regulation may be different to PAR1.

Several other studies have looked for key X chromosome genes that have lower expression in adult Turner tissues^19,20,24,27,30,41,42^. These studies have mainly analyzed blood leukocytes and more recently muscle and fat^19,30^. In addition to several PAR1 genes, the study by Viuff et al.^19^ found a subset of key genes that were differentially expressed by X chromosome dosage (45,X *versus* 46,XX and 47,XXY *versus* 46,XY), including genes that escape X inactivation (e.g., *PUDP*, *ZFX*, *KDM6A*, *JPX*, *TSIX* and *XIST*). These genes were also generally found to be differentially expressed in our dataset. Another recent study by San Roman et al.^25^ proposed a core group of X chromosome genes that are linked to sex differences. Several of these genes were found to be lower in our datasets, such as *DDX3X*, *KDM5C, KDM6A*, *SMC1A* and *ZFX.* The role of histone demethylases (e.g., *KDM5C*, *KDM6A*) may be particularly important. For example, pathogenic variants in *KDM6A* cause Kabuki syndrome 2 (KABUK 2, X-linked) (OMIM 300867), which has a high prevalence of TS-associated features such as horseshoe kidney, short stature and hearing loss^43^. Here, we report, for the first time, core groups of PAR and XCI genes that show lower expression in key biologically-relevant fetal monosomy X tissues such as pancreas, kidney, liver and skin, rather than the typical approach of studying adult samples, usually blood leukocytes.

Extending our analysis to other genes with lower expression (log_2_FC<-0.35) in monosomy X identified several autosomal genes which may be clinically relevant. For example, lower expression of the low-density lipoprotein receptor gene (*LDLR*), which regulates cholesterol homeostasis, could contribute to the risk of dyslipidemia in Turner syndrome^44^. Furthermore, pathway enrichment analysis identified biological processes related to ascending/thoracic aortic aneurysm and connective tissue disease^33,45^. Aortic root dilatation and thoracic aortic aneurysm (TAA) is rare in the population but can be a significant risk in women with TS^16,46^. Aortic root dilatation is now regularly monitored in adulthood^3,15,17,45^ and TAA has the highest odds ratio for mortality in the adult TS group (standardized mortality ratio, =23; 95% CI 13.8– 37.8)^3,47^. The main genes identified in this pathway were collagen (*COLA*) genes and fibulin 1 (*FBN1*). Defects in *FBN1* cause Marfan syndrome, which is also associated with TAA^33^. This finding raises the possibility of a pathogenic link between the TS and Marfan syndrome.

We also hypothesize that this risk of TAA could be compounded by lower expression of *AGTR2* (angiotensin II receptor type 2, AT2R) in monosomy X. *AGTR2* typically antagonizes the effects of the angiotensin II type 1 receptor (AT1R). Lower *AGTR2* could lead to higher blood pressure and vascular proinflammatory effects^48^. Several studies have shown higher plasma renin activity in women with TS^49,50^. In addition, monosomy X augments the severity of angiotensin II-induced aortopathies in mice^51^. Notably, AT1R blockade has been shown in recent systematic reviews to reduce the risk of thoracic aortic aneurysm in Marfan syndrome^52^. Further studies are needed to assess whether the interplay of these factors could lead to increased TAA risk in women with TS and be an avenue for new therapeutic approaches.

Although most of the effects of TS might be expected to be due to a reduction of gene dosage, one gene (*OVCH1-AS1*) was consistently upregulated in all datasets studied. *OVCH1-AS1* encodes a long non-coding RNA that is expressed at relatively low transcript level, which is typical for lncRNAs, but with at least two-fold increase in expression in all 45,X tissues assessed. The potential role of *OVCH1-AS1* is not known. Recent studies have suggested different expression levels of leukocyte *OVCH1-AS1* in relation to frailty with age in humans^34^, as well as higher OVCH1-AS1 expression in biopsies from Crohn’s disease^53^. Furthermore, several studies have identified *OVCH1-AS1* as a higher-expressed gene in leukocytes, fat, and muscle from adult women^19,20,24,27,41^. This finding was rarely highlighted in detail, but here we have conclusively demonstrated that *OVCH1-AS1* is higher in monosomy X in 14 different sub-studies from multiple different tissues.

As antisense lncRNAs sometimes modulate sense strand RNA, or genes in the region, we undertook more detailed analysis of this region of chromosome 12, but we did not see an obvious effect on the expression of *OVCH1* nor of other local genes. Of note, like many lncRNAs, this gene is present in humans but not in the mouse. We hypothesize that, if *OVCH1-AS1* is biologically relevant, this feature could contribute to marked differences in phenotype between monosomy X in humans and the murine model of a single X chromosome^54^.

Some of the biggest questions to address are how monosomy X is such a common cause of pregnancy loss and also whether there are any specific features in those pregnancies that survive to term? It has been estimated that 98% of monosomy pregnancies are lost mostly towards the end of the first trimester and that this contributes to a substantial proportion of spontaneous miscarriage in the general population^4,36,55^.

One long-standing hypothesis is that term monosomy X pregnancies have placental mosaicism for a fetally-derived 46,XX line, but direct data are limited^36,56^. Here, we did not identify significant 46,XX mosaicism using both RNA (*XIST* counts) and DNA (SNP arrays) technologies in the six 45,X placental samples studied, suggesting that placenta mosaicism is not likely to be a common cause of fetal rescue in TS, although placental sampling only occurred in two distinct regions.

Another question relates to the mechanistic basis of pregnancy loss. Several studies have proposed that PAR genes such as *CSF2RA* are implicated, but direct data are limited^2,6^. We therefore developed a systematic approach to identify placenta-specific genes with lower expression in monosomy X placenta compared to control placenta. We found two genes of interest: *AADACL3* and *CSF2RA*.

*AADACL3* encodes arylacetamide deacetylase-like 3, a putative membrane hydrolase expressed at low level, primarily in the extravillous trophoblast. *AADACL3* expression was more variable between monosomy X samples and controls. The functional role of this gene in the placenta is unclear, although higher *AADACL3* expression has been reported in 46,XY preterm and term placental tissue compared to 46,XX^57^.

More importantly, the PAR1 gene *CSF2RA* was observed in our dataset as having consistently lower expression in monosomy X compared to both 46,XX and 46,XY control placenta samples. In scRNA-seq analysis, *CSF2RA* is clearly expressed in early trimester and term placenta, as well as in macrophages, NK-cells and activated and resting T-cells, consistent with its expression in placenta and hematopoietic system-derived tissues in adult panels^37,38,58^. CSF2RA encodes the alpha subunit of the CSF receptor, which mediates the effects of CSF2 (GM-CSF) on hematopoietic differentiation, immunity, and inflammation.

Immune regulation within the fetal-placental interface is emerging as a key mechanism in pregnancy maintenance, and *CSF2RA* has been proposed as a candidate gene for early lethality in 45,X embryos^2,6^. *CSF2RA* has been shown to have 10-fold lower expression in 45,X-derived human embryonic stem cells compared to 46,XX-derived controls, and reduced expression in BDCA4+ dendritic cells from women with TS^6,59^. *CSF2RA* is implicated in placental development^60^, and studies in bovine cultured embryos suggest CSF2 promotes embryo survival through antiapoptotic effects and by signaling of developmental pathways^61^. However, direct analysis of monosomy X placenta has not been reported.

Here, we show morphological changes in the monosomy X placenta during development. Using IHC, we have shown that CSF2RA localizes to syncytial trophoblast and basal trophoblast cells, with more diffuse patterning in monosomy X placenta at 14wpc at a time of degeneration of the brush border. Coupled with this, pathway analysis of lower-expressed genes in the monosomy X placenta showed enrichment of key biological processes, such as leukocyte activation, immune responses, and inflammation. Taken together, these findings suggest that CSF2RA is likely to be a key modulator of fetal-placental dysfunction in monosomy X pregnancies and might have implications for our understanding of pregnancy loss in general.

This study has several limitations. Firstly, access to tissues of interest was limited so only a relatively small number of samples were available and within a relatively narrow developmental time window. However, having samples within a 4-week period reduced marked variations due to organ development, increasing the power to detect more subtle sex chromosome-related effects. Second, one fetus had low level mosaicism for a Y chromosome line. The overall effects of this variability on our findings were likely to be minimal. Finally, alternative approaches such as scRNA-seq might have provided more granular detail of single cell transcriptomics, but risked sampling bias and limited detection of low-level RNA transcripts. Thus, a bulk RNA-seq strategy was used with robust experimental design, replicating data with both 46,XX and 46,XY control groups, and validating findings in multiple different tissue groups. This is important as gene dosage changes linked to loss of a single X chromosome are generally subtle and of a lower magnitude than in most studies of developmental differential gene expression.

To the best of our knowledge, this is the first detailed transcriptomic study of human monosomy X in key clinically relevant tissues (e.g., pancreas, kidney, liver, skin, and placenta). This work has highlighted several novel findings with implications for sex chromosome biology, the underlying mechanisms that could contribute to TS and potential targeted therapeutics. These findings will be important for facilitating research in the future.

## MATERIALS AND METHODS

### Tissue samples

Human fetal tissue samples included in this study were obtained with informed consent and with ethical approval (REC references: 18/LO/0822, 18/NE/0290) from the Medical Research Council (MRC)/Wellcome Trust-funded Human Developmental Biology Resource (HDBR) (http://www.hdbr.org). The HDBR is a biobank regulated by the U.K. Human Tissue Authority (HTA) (www.hta.gov.uk). Fetal age was estimated by measuring foot-length and knee-heel length linked to standard growth data. Fetal karyotype assessment was undertaken using quantitative PCR (chromosomes 13, 15, 16, 18, 21, 22 and X and Y) and confirmed with array analysis when aneuploidy was suspected. Tissues with a monosomy X (45,X) karyotype were stored at −70°C. Control samples (46,XX, 46,XY), matched for tissue and age, were also obtained from the HDBR.

### Study design

A range of tissues, that are potentially relevant to the clinical phenotypes in Turner syndrome, were used for transcriptomic analysis. These included pancreas (diabetes mellitus), liver (liver enzyme elevation), kidney (developmental anomalies, renal dysfunction) and skin (naevi), as well as a “mixed group” that was comprised of brain, spleen, lung, and heart (Fig. 1a). Each sequenced group consisted of four monosomy 45,X samples, with four age and tissue matched 46,XX control samples and four age and tissue matched 46,XY control samples, between 11 and 15 wpc. For the 45,X pancreas group, a 13 wpc sample was bisected sagittally from the head of the pancreas to the tail, so that RNA could be independently extracted twice from the same biological sample. Each organ in the mixed tissue group (brain, spleen, lung, and heart) was collected at a different age stage within the timeframe (11 to 15 wpc) (Fig. 1a). An addition group of muscle samples (n=4 each group) was included for validation of *OVCH1-AS1* expression.

For the placental transcriptomic study, six monosomy 45,X placenta samples were obtained together with six age matched 46,XX control samples and six age matched 46,XY control samples, between 11 and 15wpc. Samples were stored in −70°C. Additional second samples of tissue (−70°C) were taken from a different region of the 45,X placenta for the assessment of potential mosaicism. Independent placental samples were stored in 10% formalin for histology and immunohistochemistry.

### RNA/DNA extraction

Samples were removed from −70°C storage and dissected further, where necessary, using a standardized approach so that the required amount of tissue was available (up to 30 mg). RNA and DNA extractions were performed using an AllPrep DNA/RNA Mini Kit following the manufacturer’s protocol (Qiagen, Hilden, Germany). Tissues were immediately cut into 1mm^3^ pieces on ice and transferred to lysis buffer supplied (buffer RLT). Samples were dissociated in this lysis buffer using a Kimble™ Kontes™ motorized pellet pestle (DWK Life sciences, Mainz Germany). To isolate RNA, the first flow through from the AllPrep column was digested using proteinase K, subsequent ethanol washes undertaken to allow binding of total RNA including miRNA to the RNeasy mini spin column, followed by Dnase I digestion and further washes to ensure elution of high-quality RNA. RNA quality was assessed using a Tapestation 4200 platform (Agilent Technologies, Santa Clara, CA, USA). DNA was also extracted from the same samples using the manufacturer’s protocol. In brief, homogenized lysate was passed through an AllPrep DNA mini spin column to bind genomic DNA. Following proteinase K digestion under optimised buffer conditions, the column was washed, and DNA eluted and quantified using NanoDrop spectrophotometer (Thermo Fisher Scientific, Waltham, MA, USA).

### SNP array analysis

In order to determine sex chromosome complement and potential mosaicism in all samples, SNP array analysis was undertaken on extracted DNA samples, The Illumina Global Screening Array platform was used (v3.0) (Infinium HTS Assay Reference Guide (# 15045738 v04) (Illumina, Inc. San Diego, CA, USA). Output data were analyzed in Illumina Genome Studio version 2.0. X chromosome mosaicism was assessed using methods described by Conlin et al., 2010^62^.

### Bulk RNA library preparation and sequencing

For bulk RNA-seq studies, RNA was used to generate cDNA libraries using a KAPA mRNA HyperPrep Kit (Roche, Basel, Switzerland) on a Hamilton Starlet robotic platform (Hamilton Company, Reno, NV, USA) and quality was analyzed using a Tapestation 4200 platform (Agilent Technologies, Santa Clara, CA, USA). Libraries were subsequently sequenced on a NovaSeq S2 Flowcell (paired end 2×56bp) (Illumina). All samples in this study were prepared and sequenced at the same time, to avoid potential batch effects. Fastq files were processed by FastQC and aligned to the human genome (Ensembl, GRCh 38.86) using STAR (v2.7.3a). The matrix containing uniquely mapped read counts was generated using featureCounts part of the R package Subread.

### Bulk RNA-seq data analysis and data representation

The following analysis was performed in R (version 4.2.2). Comparison of RNA-seq sample dimensionality was undertaken for all samples and for each individual tissue group using principal component analysis (PCA) and plots were generated using ggplot2^63,64^. Pairwise differential-expression analysis of tissues related to karyotype was performed using DESeq2^65^, comparing either four 45,X samples with four 46,XX samples, or four 45,X samples with four 46,XY samples. For the placental study, six samples were used in each group. Data were generated as log_2_ fold change (FC) differences between groups >0.5 and were considered statistically significant with an adjusted p-value <0.05. Volcano plots were generated using ggplot2. Heatmaps for differentially expressed genes in 45,X tissue compared to control 46,XX and 46,XY controls were generated using the pheatmap library in R. Violin plots of normalized counts of tissues were plotted in ggplot2 in R. Venn diagram analyses between different subsets of expressed genes was undertaken using Venny 2.1 or InteractiVenn^66,67^. Pathway analysis and enrichment analysis was undertaken using clusterProfiler^68^, Metascape^69^ and gProfiler^70^ version: e110_eg57_p18_4b54a898.

### OVCH1-AS1 gene locus and conservancy

The *OVCH1-AS1* (*OVCH1 Antisense RNA 1*) gene locus (including neighboring genes) was defined using the University of California, Santa Cruz (UCSC) Genome Browser (human GRCh38/hg38; Chromosome12:29,389,642-29,487,473) and the Ensembl browser 110 (Human GRCh38.p13; Chromosome 12: 29,387,326-29,489,451). Conservation of the *OVCH1-AS1* locus was evaluated in the UCSC genome browser (Multiz alignments of 100 vertebrates) using default species settings (Chimp, Rhesus monkey, Mouse, Dog, Elephant, Chicken, *Xenopus tropicalis* and Zebrafish).

### Quantitative real-time PCR (qRT-PCR)

cDNA was generated from pancreas, liver and kidney RNA using SuperScript III reverse transcriptase (Thermo Fisher Scientific Inc, MA, USA). Quantitative-real-time polymerase chain reaction (qRT-PCR) was performed using TaqMan Fast Advanced MasterMix (Applied Biosystems MA, USA) and Taqman gene expression assays for *OVCH1-AS1* (Hs04333030_m1) and genes in the surrounding locus (*OVCH1* (Hs07289759_m1)*, ERGIC2* (Hs00275449_m1)*, FAR2* (Hs00216461_m1)*, TMTC1* (Hs00405786_m1)) (Applied Biosystems, MA, USA). Analysis was carried out using the comparative CT (2^-ΔΔCT^) method^71^. *ACTB* (Hs01060665_g1) or *GAPDH* (Hs02786624_g1) were used as a housekeeping genes. Data were normalized to 13wpc 46,XY samples for each organ. The gene *OVCH1-AS1* was assessed in triplicate on three independent occasions and mean 2^-ΔΔCT^ was obtained for each study. Genes in the region of *OVCH1-AS1* (*OVCH1*, *ERGIC2*, *FAR2* and *TMTC1*) were assessed in triplicate on one occasion. Data were plotted using GraphPad Prism (version 9.5 for Windows, GraphPad Software, San Diego, CA, USA, www.graphpad.com). Differences between groups were analyzed using one-way ANOVA (Kruskal-Wallis) (GraphPad Prism).

### In silico assessment of OVCH1-AS1

A systematic review of publications citing *OVCH1-AS1* was undertaken in PubMed and GoogleScholar, using the gene name as a search term (October 2023). Although *OVCH1-AS1* is considered a long non-coding RNA (lncRNA) (GeneCards GC12P029389, Ensembl ENSG00000257599), the potential to generate a protein coding transcript was analyzed using the following algorithms: CPAT coding probability, PhyloCSF score, PRIDE reprocessing 2.0, Lee translation initiation sites, Bazzini small ORFs (accessed via RNAcentral and Lncipedia, October 2023). Gene expression of OVCH1-AS1 and exon usage in adult tissues was obtained from GTEx (https://www.gtexportal.org/home/gene/OVCH1-AS1; ENSG00000257599.1; exon usage Source symbol;Acc:HGNC:44484).

### Placenta immunohistochemistry (IHC)

Placental samples for staining were fixed in 10% formalin before being processed, wax embedded and sectioned (3 µm). Hematoxylin and eosin (H&E) stains were performed using standard protocols. IHC for CSF2RA (Colony Stimulating Factor 2 Receptor Subunit Alpha) was undertaken on 3µm sections using a Leica Bond-max automated platform (Leica Biosystems). In brief, antigen retrieval was performed to unmask the epitope (Heat Induced Epitope Retrieval (HIER), Bond-max protocol F), endogenous activity was blocked with peroxidase using a Bond polymer refine kit (cat # DS9800), then incubation was undertaken with a primary rabbit polyclonal CSF2RA (GM-CSF receptor alpha) antibody for 1 hour (Origene TA323990S, 1:100 dilution, HIER1 for 30 mins). A post-primary antibody was applied to the sections (Bond polymer refine kit) and horseradish peroxidase (HRP) labeled polymer, followed by 3, 3-diaminobenzidine (DAB) chromogen solution. Sections were counterstained with hematoxylin, washed, dehydrated, cleared in two xylene changes and mounted for light microscopy. Images were taken on an Aperio CS2 Scanner (Leica Biosystems) at 40x objective. Analysis was performed with QuPath (v.0.2.3) (https://qupath.github.io) software.

### Tissue and placental expression of key genes

General expression of *AACAD3L* and *CSF2RA* in adult tissues was performed using GTEx Consensus bulk RNA-seq consensus data. Single-cell RNA-seq expression of these genes in first trimester placenta was visualized using the CELLxGENE^39^ repository to access data by Vento-Tormo et al.,2018^37^ (https://maternal-fetal-interface.cellgeni.sanger.ac.uk/) (CC-BY-4.0). Single-cell RNA-seq expression of these genes in term and pre-term placenta was visualized using data by Pique-Regi et al.,2019^38^ (http://placenta.grid.wayne.edu/) (CC-BY-4,0). Consensus tissue datasets were obtained from the Human Protein Atlas for *AADACL3* expression (https://www.proteinatlas.org/ENSG00000188984-AADACL3/tissue) and *CSF2RA* expression (https://www.proteinatlas.org/ENSG00000198223-CSF2RA/tissue) (Data accessed 10/17/23; Protein Atlas version 23.0) (Uhlén et al.,2015)^58^ (CC-BY-4.0)

### Statistical analyses

Statistical analysis for quantitative variability in relative expression between karyotypes was undertaken using one-way ANOVA (Kruskal-Wallis) test in GraphPad Prism and presented using GraphPad (version 9.5.1 for Windows, GraphPad Software, San Diego, CA, USA, (www.graphpad.com). Statistical significance associated with differentially expressed genes were carried out in R, as defined above^63,64^. A p-value of <0.05 was considered significant, following adjustment for multiple comparisons where indicated.

## Contributors

JPS and JCA conceptualized the study. JPS, IDV, FB, SMB, TB, OO, DL, ND, GMK, KM, LN, ARM, MI, NM, GEM, BC, NS, GSC and JCA undertook data curation. JPS and SMB undertook formal data analysis. JCA was involved in funding acquisition. JCA and NS oversaw project administration and supervision. IDV undertook validation of bioinformatic platforms and pipelines. JPS and FB were responsible for data visualization. JPS and JCA wrote the original draft with input from GSC. All authors were involved in reviewing and editing the final manuscript. JPS and JCA had full access to all data in the study and had final responsibility for the decision to submit for publication.

## Acknowledgments

This research was funded in whole, or in part, by the Wellcome Trust (J.C.A. 209328/Z/17/Z; S.M. McB 216362/Z/19/Z). For the purpose of Open Access, the author has applied a CC BY public copyright license to any Author Accepted Manuscript version arising from this submission. Human fetal material was provided by the Joint MRC/Wellcome Trust (Grant MR/R006237/1, MR/X008304/1 and 226202/Z/22/Z) Human Developmental Biology Resource (http://www.hdbr.org). All research at UCL Great Ormond Street Institute of Child Health is made possible by the NIHR Great Ormond Street Hospital Biomedical Research Centre (grant IS-BRC-1215-20012). The views expressed are those of the authors and not necessarily those of the National Health Service, National Institute for Health Research, or Department of Health.

The Genotype-Tissue Expression (GTEx) Project was supported by the Common Fund of the Office of the Director of the National Institutes of Health, and by NCI, NHGRI, NHLBI, NIDA, NIMH, and NINDS. The data used for the analyses described in this manuscript were obtained from the GTEx Portal on 10/10/23.

## Data availability

Bulk RNA-sequencing data are deposited in ArrayExpress/Biostudies (accession number E-MTAB-13673).

**Supporting data** (Fig3b to Fig.7h; Supplementary Fig 2a to 10)

**Supplementary Information** (Supplementary Table 1; Supplementary Figures 1-15)

Supplementary Table 1. OVCH1-AS1 protein coding potential

Supplementary Figure 1. Extended experimental design

Supplementary Figure 2. Overview of Y gene counts in all samples studied and array analysis in a 45,X fetus with a Y chromosome line

Supplementary Figure 3. Principal component analysis (PCA) of tissues

Supplementary Figure 4. Genes with lower expression in monosomy X samples

Supplementary Figure 5. Pseudoautosomal region 1 (PAR1) gene expression in different tissues and for different karyotypes

Supplementary Figure 6. Pseudoautosomal region 2 (PAR2) gene expression in different tissues and for different karyotypes

Supplementary Figure 7. Genes with higher expression in monosomy X samples

Supplementary Figure 8. OVCH1-AS1 expression in the Genotype-Tissue Expression (GTEx) dataset

Supplementary Figure 9. Expression of additional genes in the OVCH1-AS1 locus

Supplementary Figure 10. Placenta study design and XIST counts in placental samples and placental biological replicates

Supplementary Figure 11. SNP arrays from monosomy X and control placentae for the X chromosome

Supplementary Figure 12. Principal component analysis (PCA) of 46,XX and 46,XY placental samples and control tissue samples

Supplementary Figure 13. AADACL3 and CSF2RA expression in the Human Protein Atlas Consensus dataset

Supplementary Figure 14. Placental single cell RNA-sequencing expression of AADACL3 and CSF2RA in first trimester placenta

Supplementary Figure 15. Placental single cell RNA-sequencing expression of AADACL3 and CSF2RA at parturition

**Supplementary Data** (files 1-41)

Supplementary Data 1: Samples used in this study

Supplementary Data 2: Pancreas tissue group comparison of 45,X vs 46,XX - higher in monosomy X

Supplementary Data 3: Pancreas tissue group comparison of 45,X vs 46,XX - lower in monosomy X

Supplementary Data 4: Pancreas tissue group comparison of 45,X vs 46,XY – higher in monosomy X

Supplementary Data 5: Pancreas tissue group comparison of 45,X vs 46,XY – lower in monosomy X

Supplementary Data 6: Liver tissue group comparison of 45,X vs 46,XX – higher in monosomy X

Supplementary Data 7: Liver tissue group comparison of 45,X vs 46,XX – lower in monosomy X

Supplementary Data 8: Liver tissue group comparison of 45,X vs 46,XY - higher in monosomy X

Supplementary Data 9: Liver tissue group comparison of 45,X vs 46,XY - lower in monosomy X

Supplementary Data 10: Kidney tissue group comparison of 45,X vs 46,XX - higher in monosomy X

Supplementary Data 11: Kidney tissue group comparison of 45,X vs 46,XX - lower in monosomy X

Supplementary Data 12: Kidney tissue group comparison of 45,X vs 46,XY - higher in monosomy X

Supplementary Data 13: Kidney tissue group comparison of 45,X vs 46,XY - lower in monosomy X

Supplementary Data 14: Skin tissue group comparison of 45,X vs 46,XX - higher in monosomy X

Supplementary Data 15: Skin tissue group comparison of 45,X vs 46,XX - lower in monosomy X

Supplementary Data 16: Skin tissue group comparison of 45,X vs 46,XY - higher in monosomy X

Supplementary Data 17: Skin tissue group comparison of 45,X vs 46,XY - lower in monosomy X

Supplementary Data 18: Mixed tissue group comparison of 45,X vs 46,XX - higher in monosomy X

Supplementary Data 19: Mixed tissue group comparison of 45,X vs 46,XX - lower in monosomy X

Supplementary Data 20: Mixed tissue group comparison of 45,X vs 46,XY - higher in monosomy X

Supplementary Data 21: Mixed tissue group comparison of 45,X vs 46,XY - lower in monosomy X

Supplementary Data 22: Muscle tissue group comparison of 45,X vs 46,XX - higher in monosomy X

Supplementary Data 23: Muscle tissue group comparison of 45,X vs 46,XX - lower in monosomy X

Supplementary Data 24: Muscle tissue group comparison of 45,X vs 46,XY - higher in monosomy X

Supplementary Data 25: Muscle tissue group comparison of 45,X vs 46,XY - lower in monosomy

Supplementary Data 26: Pseudoautosomal Region 1 (PAR1) gene expression for all tissues

Supplementary Data 27: Pseudoautosomal Region 2 (PAR2) gene expression for all tissues

Supplementary Data 28: PAR gene expression in different tissues and karyotype comparisons, and mean log2FC

Supplementary Data 29: Relative expression of key genes identified by Viuff et al. 2023

Supplementary Data 30: Relative expression of key genes identified by San Roman et al. 2023

Supplementary Data 31: Differentially expressed gene with lower expression in monosomy X

Supplementary Data 32: Differentially expressed genes with higher expression in monosomy X

Supplementary Data 33: Expression of OVCH1-AS1 in all tissues

Supplementary Data 34: Expression of genes in the OVCH1-AS1 locus in all tissues

Supplementary Data 35: Placenta tissue group comparison of 45,X vs 46,XX - higher in monosomy X

Supplementary Data 36: Placenta tissue group comparison of 45,X vs 46,XX - lower in monosomy X

Supplementary Data 37: Placenta tissue group comparison of 45,X vs 46,XY - higher in monosomy X

Supplementary Data 38: Placenta tissue group comparison of 45,X vs 46,XY - lower in monosomy X

Supplementary Data 39: Placenta tissue group comparison of 46,XX placenta vs 46,XX control tissues - placenta specific genes

Supplementary Data 40: Placenta tissue group comparison of 46,XY placenta vs 46,XY control tissues - placenta specific genes

Supplementary Data 41: CSF2RA and AADACL3 expression in placenta

