## Supplementary Information for "The transcriptomic landscape of monosomy X (45,X) during early human fetal and placental development"

**Supplementary Table 1. *OVCH1-AS1* protein coding potential**

Protein coding potential of *OVCH1-AS1* as obtained from Incipedia using the prediction models shown.

| Metric | Raw result | Interpretation |
|---|---|---|
| PRIDE reprocessing 2.0 | 0 | non-coding |
| Lee translation initiation sites | 0 | non-coding |
| PhyloCSF score | -156.6865 | non-coding |
| CPAT coding probability | 52.02% | coding |
| Bazzini small ORFs | 0 | non-coding |

### Supplementary Figure 1

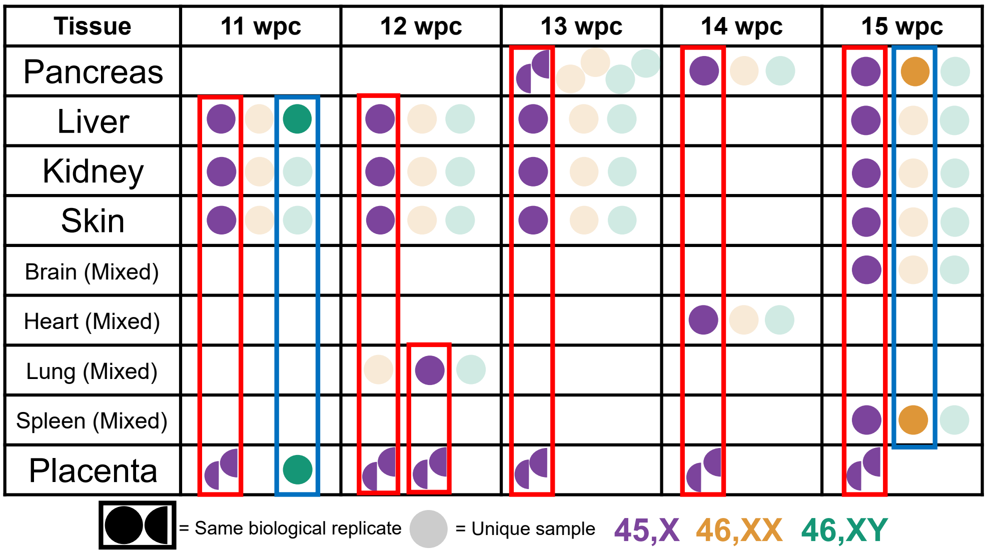


**Supplementary Figure 1. Extended experimental design.** Overview of samples used for the study, including placental samples. Those highlighted in bolder colors and in boxes refer to tissues collected from the same fetus. Lighter shaded circles indicate samples collected from different fetuses. Red boxes indicate monosomy X samples and blue boxes indicate control samples. wpc, weeks post conception.

### Supplementary Figure 2

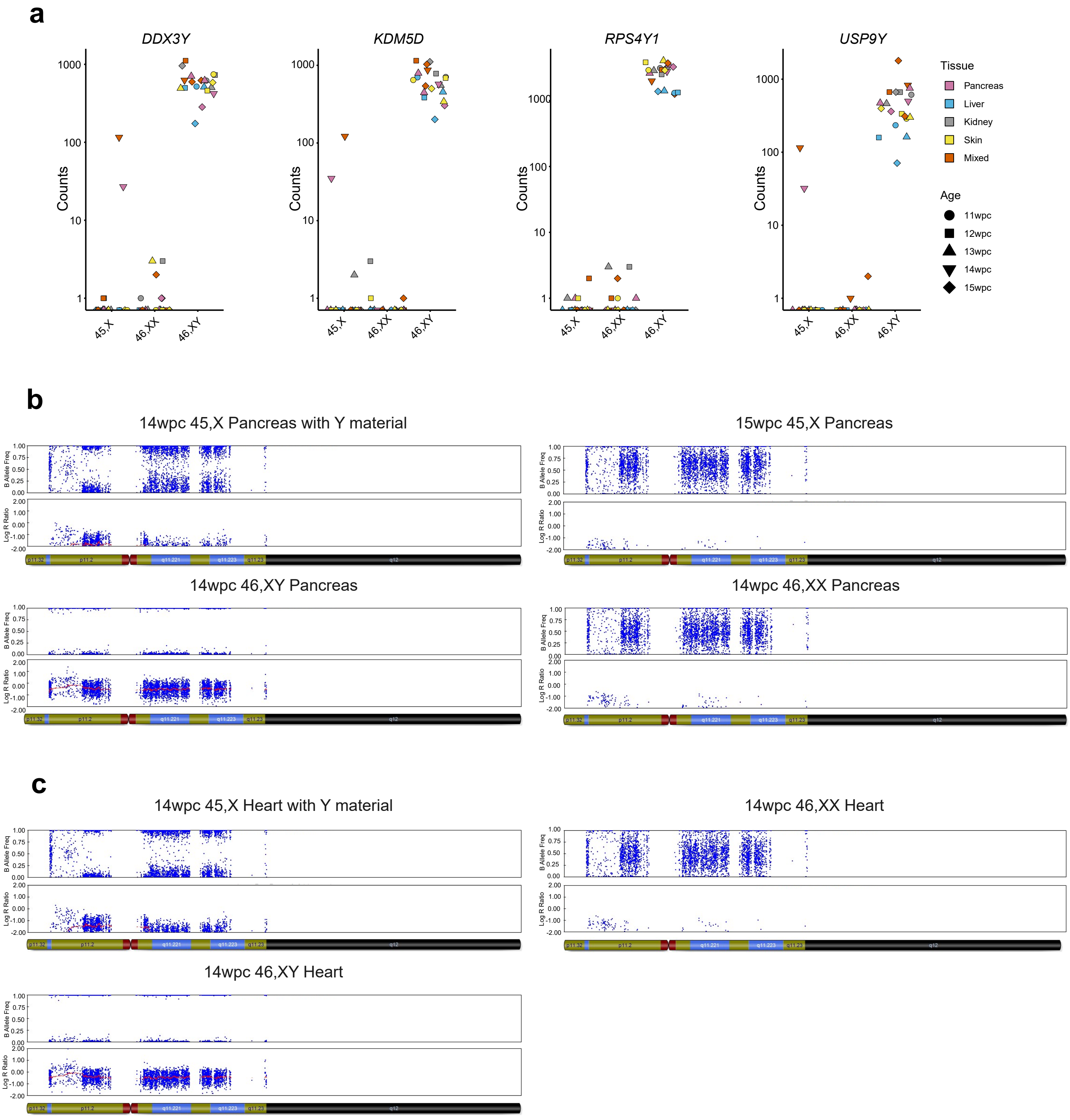


**Supplementary Figure 2. Overview of Y gene counts in all samples studied and array analysis in a 45,X fetus with a Y chromosome line. a** Raw counts from bulk RNA sequencing of selected Y chromosome genes in all tissues used for the study. Higher expression is observed in pancreas and heart samples from one 45,X fetus, suggestive of a low-level mosaic 46,XY line. Counts are shown on a log10 scale. **b** SNP array plots of the Y chromosome showing the pancreas sample with low-level Y chromosome material, and pancreas control samples for comparison (45,X, 46,XY and 46,XX). The upper panel shows the B allele frequency, which is the normalized measure of the allelic intensity ratios of two alleles A, B. The lower panel shows the log R ratio, which is the normalized measure of signal intensity for each SNP marker, as log2 of the ratio between observed and expected for two copies of the genome. The mean value is shown by the smoothened red line indicating the mean value of the Y chromosome at that position. **c** SNP array plots showing the heart sample with low-level Y chromosome material, and heart control samples for comparison (46,XY, 46,XX). No 45,X sample was included as heart was part of a mixed group. wpc, weeks post conception.

Supplementary Figure 3

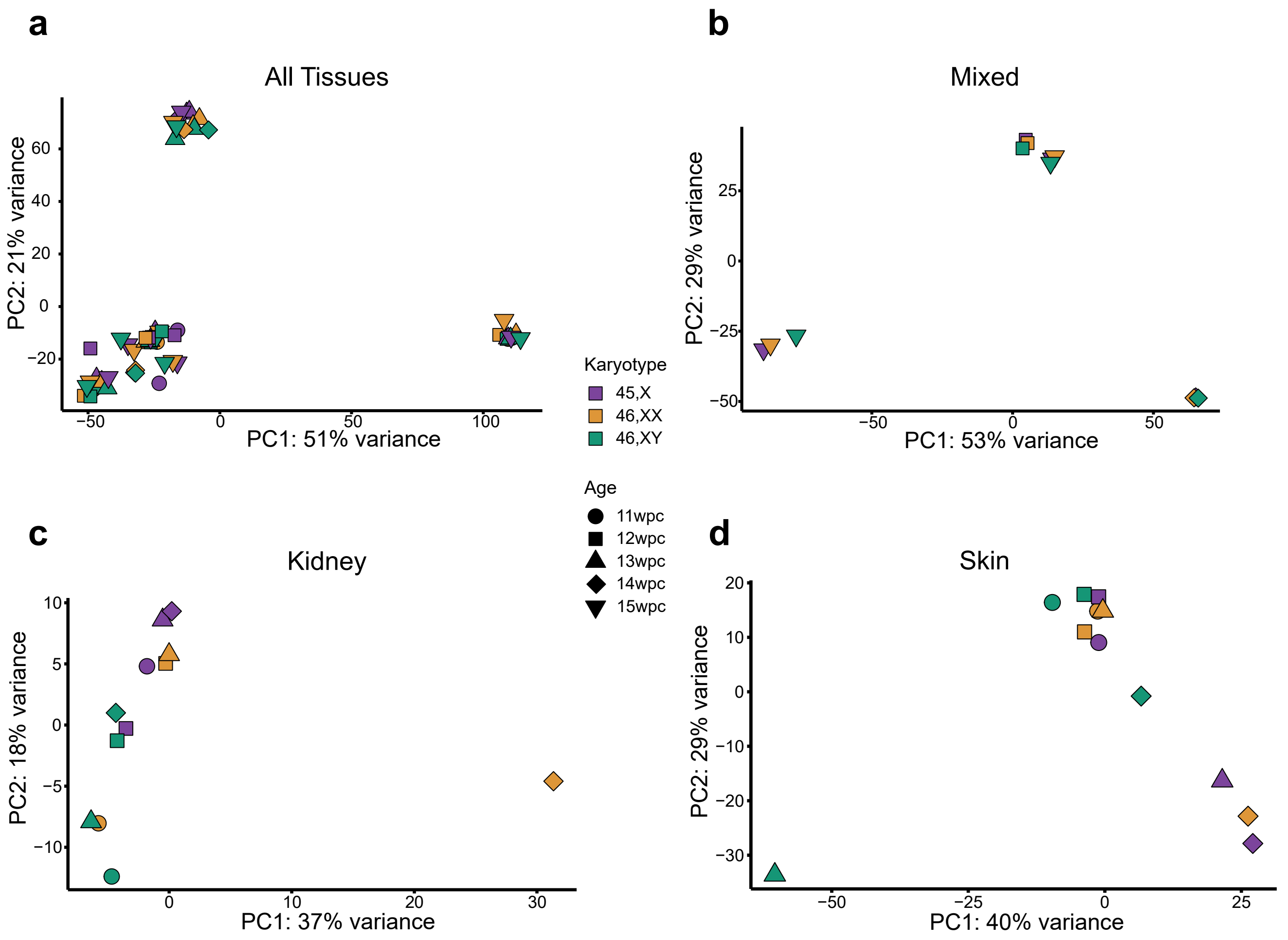


**Supplementary Figure 3. Principal component analysis (PCA) of tissues. a** PCA of all tissues colored according to karyotype. **b** PCA of mixed group colored according to karyotype. **c** PCA of all 12 kidney samples, including the one outlier (14wpc, 46,XX) with low level adrenal contamination. **d** PCA of all 12 skin samples, including the one outlier (13wpc, 46,XY) with low level muscle contamination. PC, principal component; wpc, weeks post conception.

Supplementary Figure 4

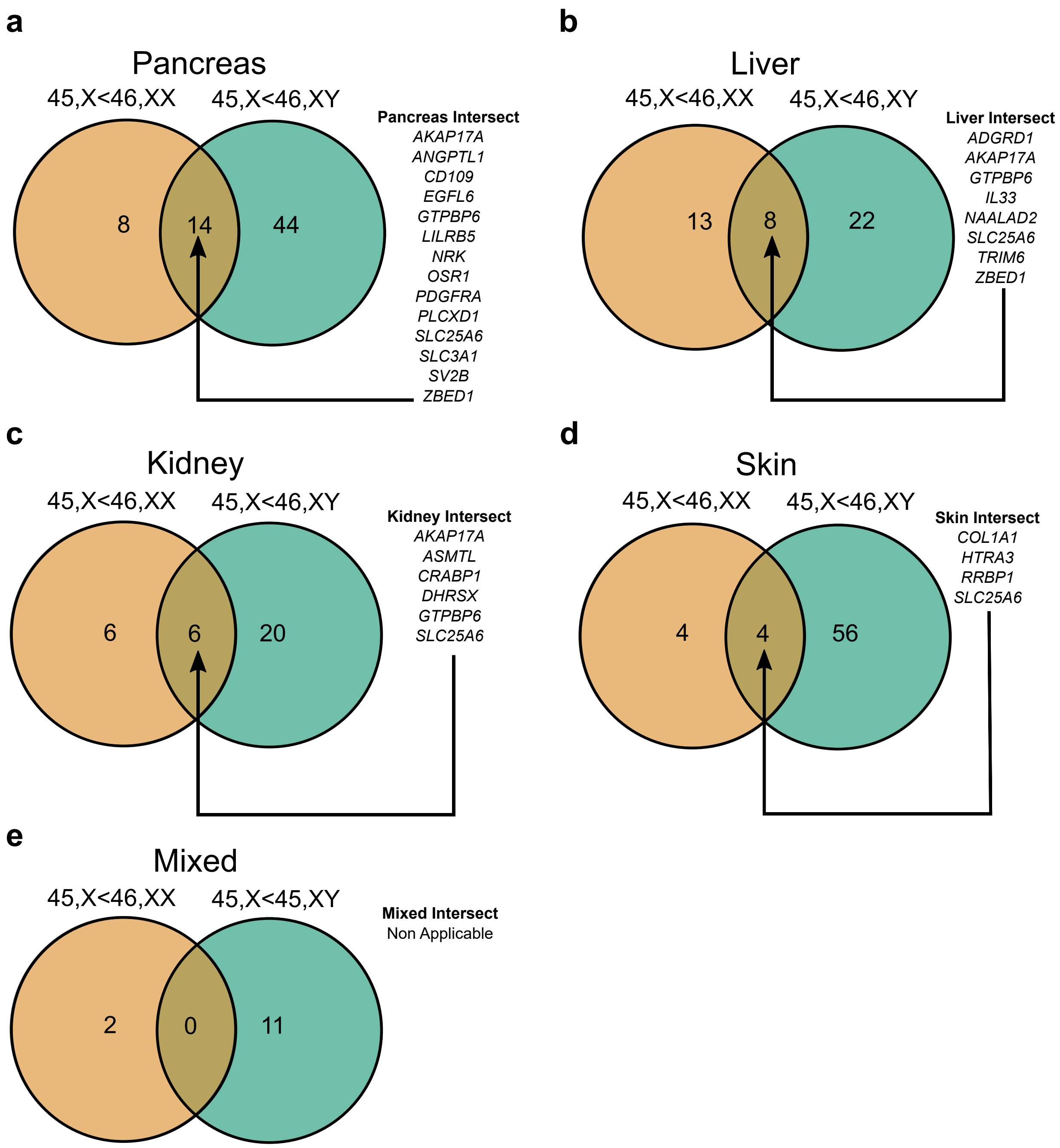


**Supplementary Figure 4. Genes with lower expression in monosomy X samples.** Overview of Venn diagram comparisons for all tissue groups studied: **a** Pancreas; **b** Liver; **c** Kidney; **d** Skin; **e** Mixed. For each 45,X *versus* 46,XX tissue study group, n=4; for each 45,X *versus* 46,XY tissue study group, n=4. Genes included where $\log_2$ fold change <-0.5; adjusted p-value <0.05.

### Supplementary Figure 5

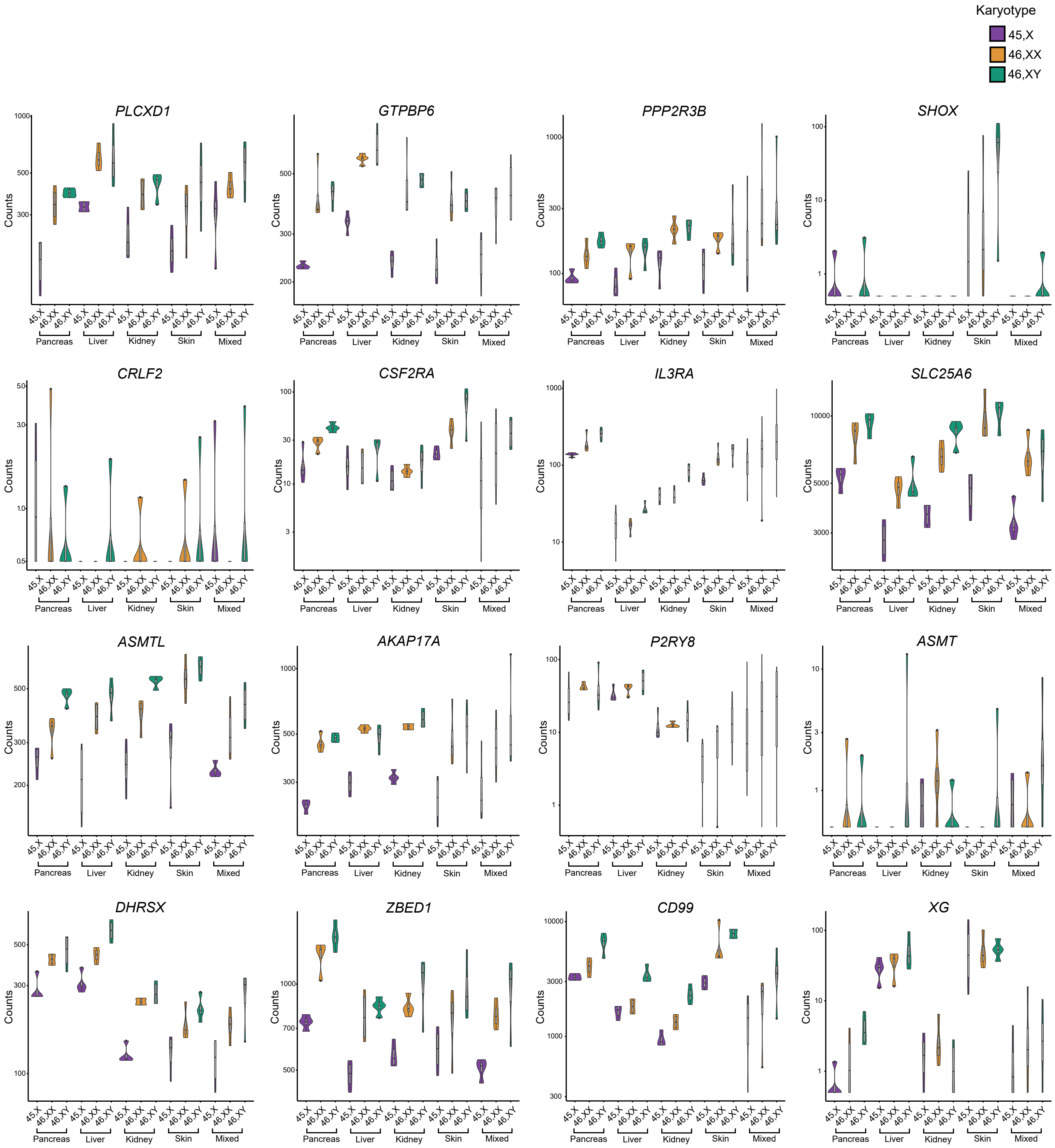


**Supplementary Figure 5. Pseudoautosomal region 1 (PAR1) gene expression in different tissues and for different karyotypes.** Violin plots of PAR1 gene normalized counts in all tissue groups studied: pancreas, liver, kidney, skin and mixed group. The karyotype order is 45,X; 46,XX and 46,XY, as indicated. Note the $\log_{10}$ scale, and considerable differences in gene expression counts between different genes.

### Supplementary Figure 6

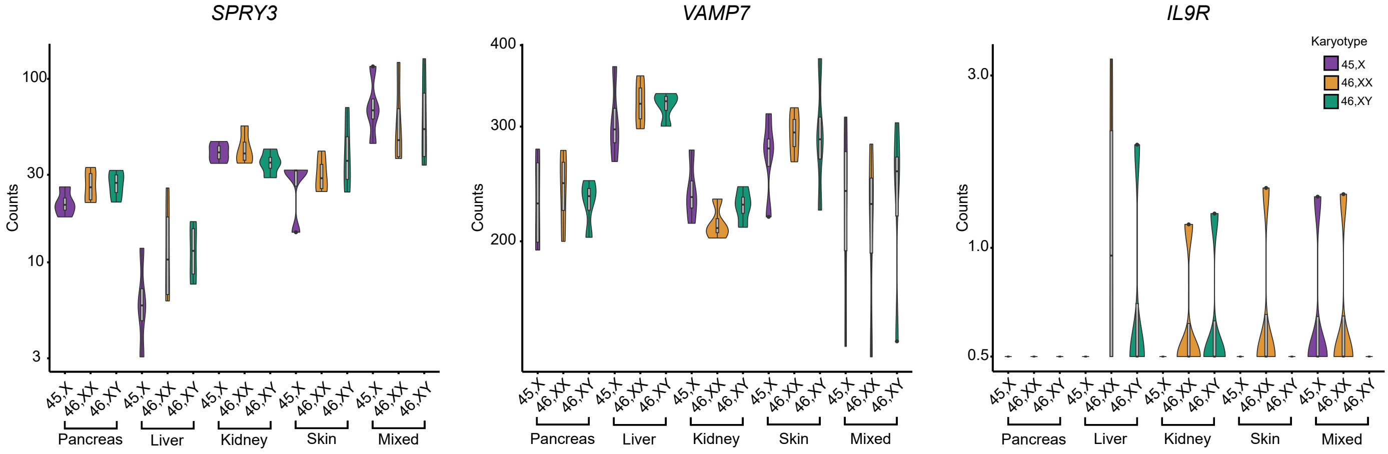


**Supplementary Figure 6. Pseudoautosomal region 2 (PAR2) gene expression in different tissues and for different karyotypes.** Violin plots of PAR2 gene counts in all tissue groups studied: pancreas, liver, kidney, skin and mixed group. The karyotype order is 45,X, 46,XX and 46,XY, as indicated. Note the $\log_{10}$ scale, and considerable differences in gene expression counts between different genes

Supplementary Figure 7

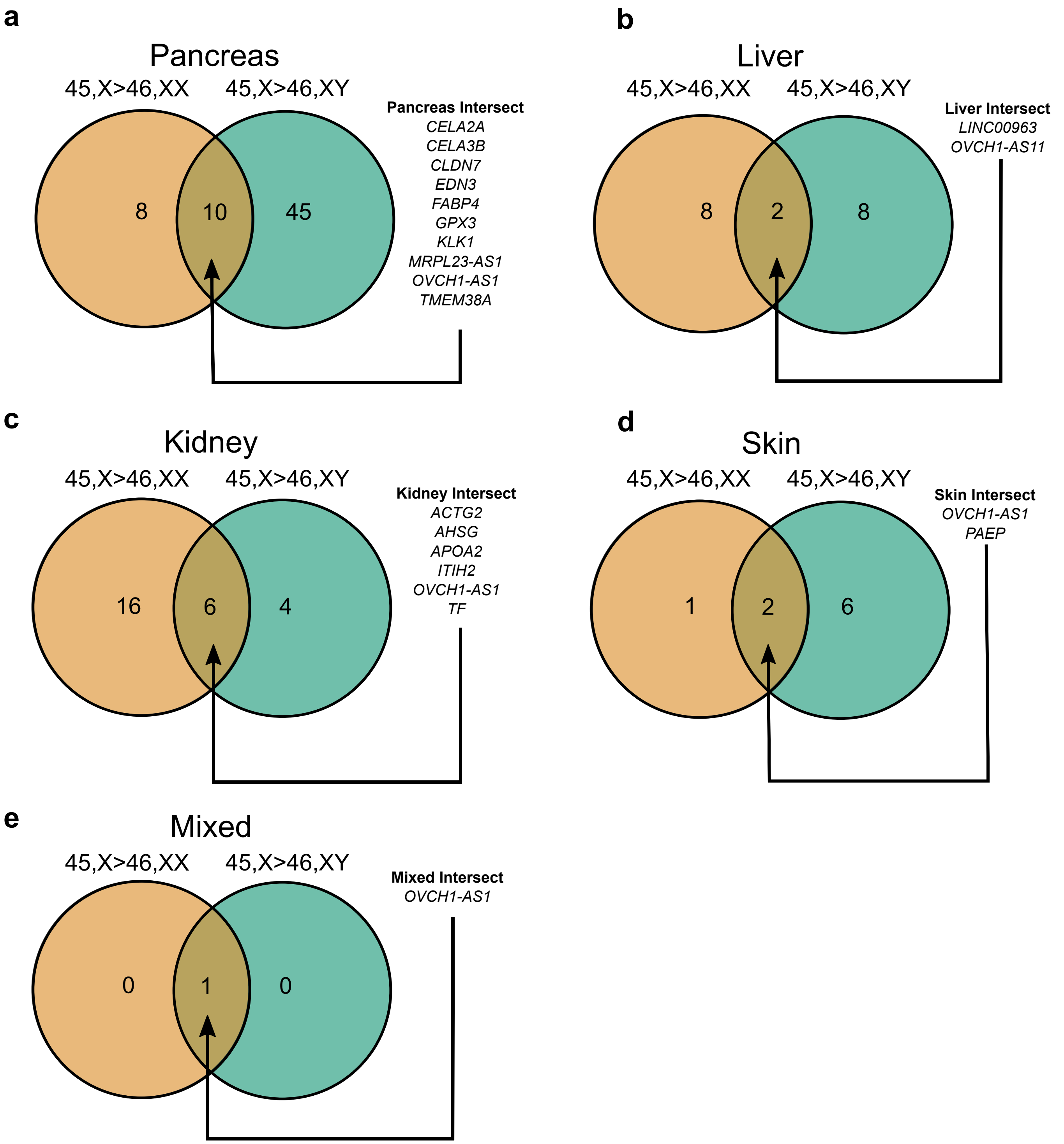


**Supplementary Figure 7. Genes with higher expression in monosomy X samples.** Overview of Venn diagram comparisons for all tissue groups studied: **a** Pancreas; **b** Liver; **c** Kidney; **d** Skin, **e** Mixed. For each 45,X *versus* 46,XX tissue study group, n=4; for each 45,X *versus* 46,XY tissue study group, n=4. Genes included where $\log_2$ fold change >0.5; adjusted p-value <0.05.

### Supplementary Figure 8

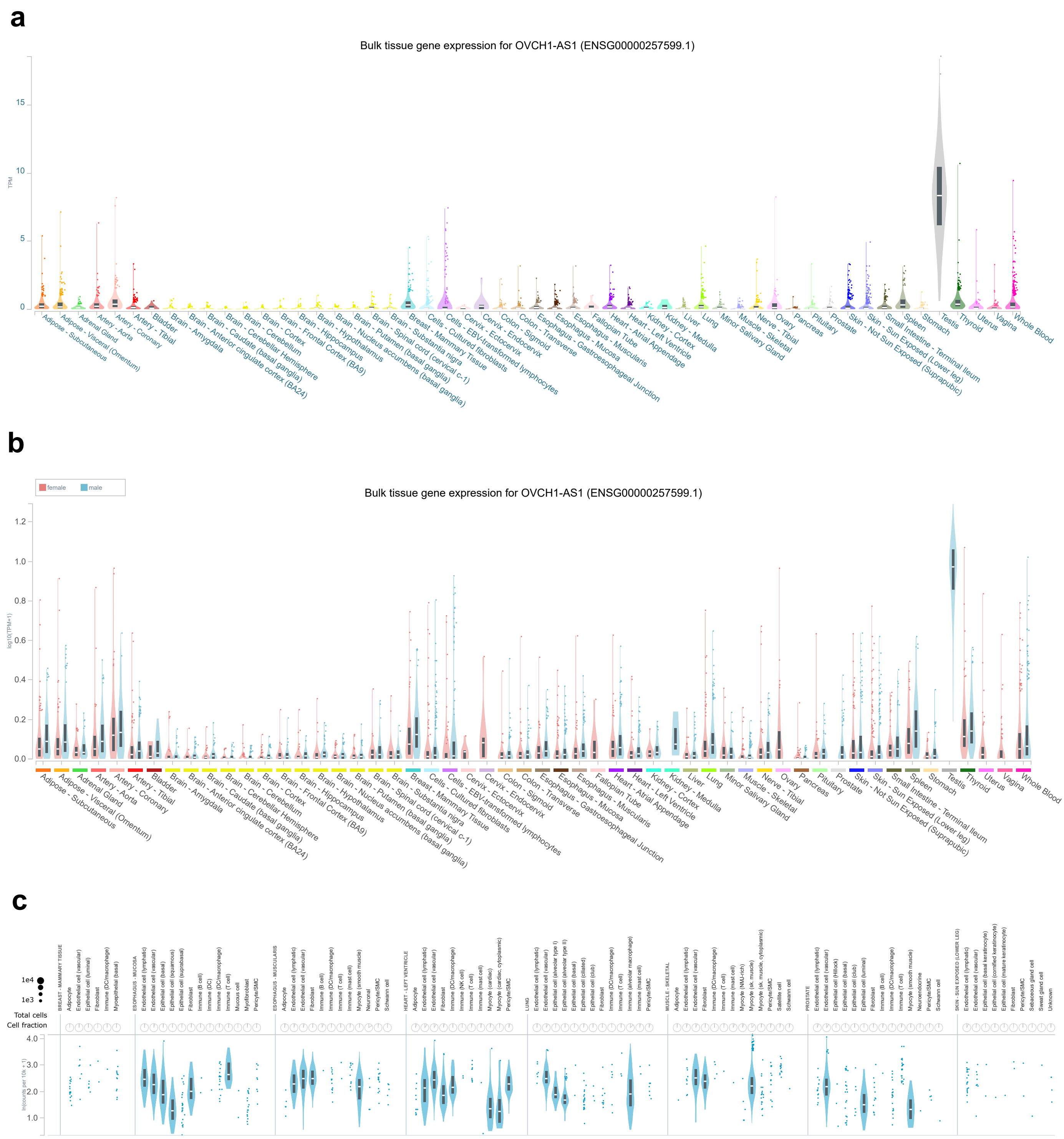


**Supplementary Figure 8. *OVCH1-AS1* expression in the Genotype-Tissue Expression (GTEx) dataset. a** Bulk RNA-seq expression for *OVCH1-AS1* in different adult tissues. **b** Bulk RNA-seq expression for *OVCH1-AS1* in different adult tissues shown by sex. **c** Single nucleus RNA sequencing (snRNA-seq) pilot data from GTEx, showing nonzero data, grouped by tissue. The Genotype-Tissue Expression (GTEx) Project was supported by the Common Fund of the Office of the Director of the National Institutes of Health, and by NCI, NHGRI, NHLBI, NIDA, NIMH, and NINDS. The data used for the analyses described in this manuscript were obtained from: [https://www.gtexportal.org/home/gene/OVCH1-AS1] the GTEx Portal on 10/17/23 (https://gtexportal.org/home) (CC-BY-4.0).

### Supplementary Figure 9

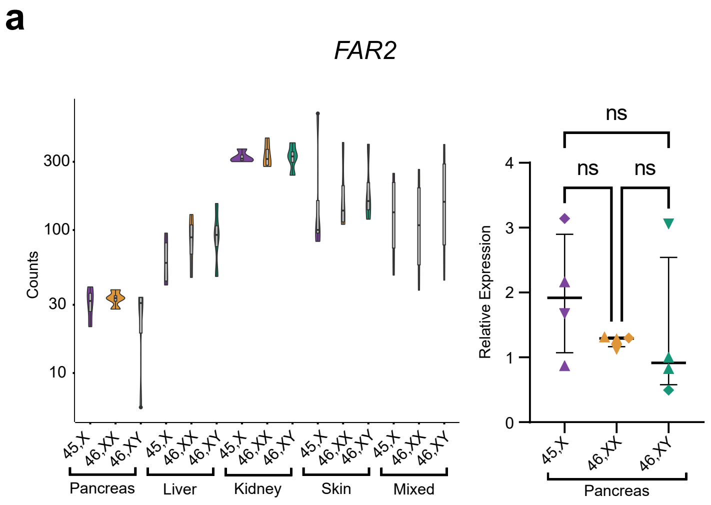


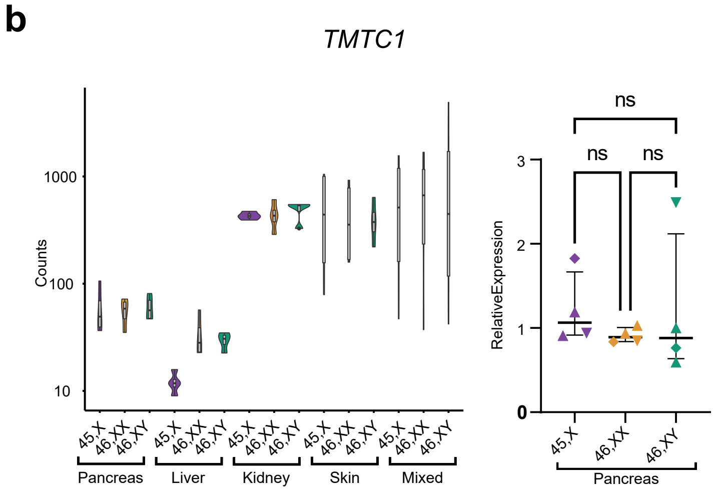


**Supplementary Figure 9. Expression of additional genes in the *OVCH1-AS1* locus. a** Violin plot of FAR2 expression (normalized counts) in all tissues (bulk RNA-seq) (n=4 each group) and quantitative real-time polymerase chain reaction (qRT-PCR) expression of FAR2 in fetal pancreas. **b** Violin plot of TMTC1 expression (normalized counts) in all tissues (bulk RNA-seq) (n=4 each group) and qRT-PCR expression in fetal pancreas. Note the $\log_{10}$ scale. ns, not significant.

### Supplementary Figure 10

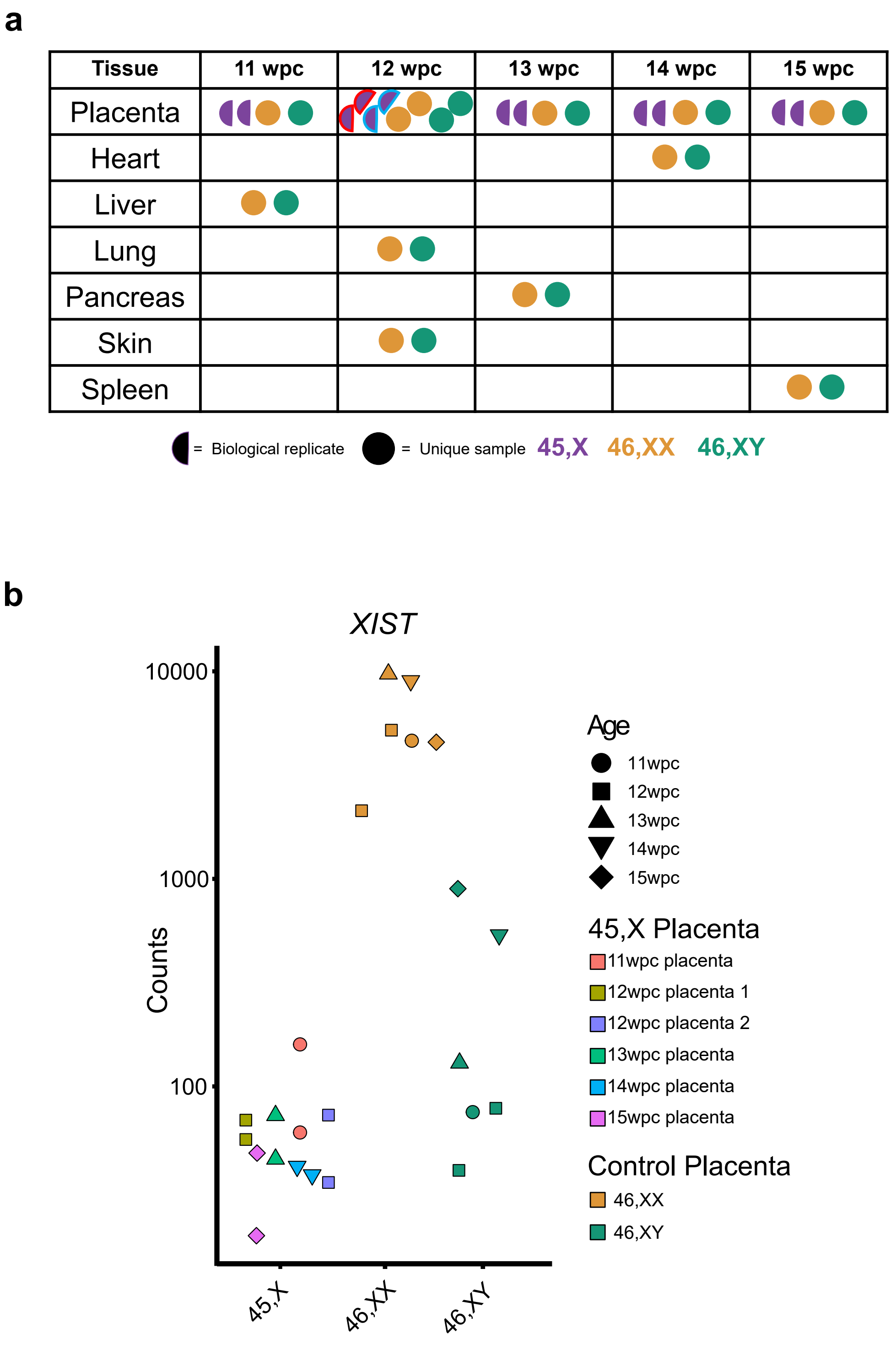


**Supplementary Figure 10. Placenta study design and *XIST* counts in placental samples and placental biological replicates. a** Overview of samples used for the placental study. Independent samples were taken from each monosomy X placenta collected. Note: two monosomy X placentae and matched controls were collected at 12 weeks post conception (red and blue outlines). **b** *XIST* raw counts for placental samples from 45,X, 46,XX and 46,XY fetuses. The two independent samples (biological replicates) from each 45,X placenta are color coded as shown in the key. Counts are shown on a $\log_{10}$ scale.

Supplementary Figure 11

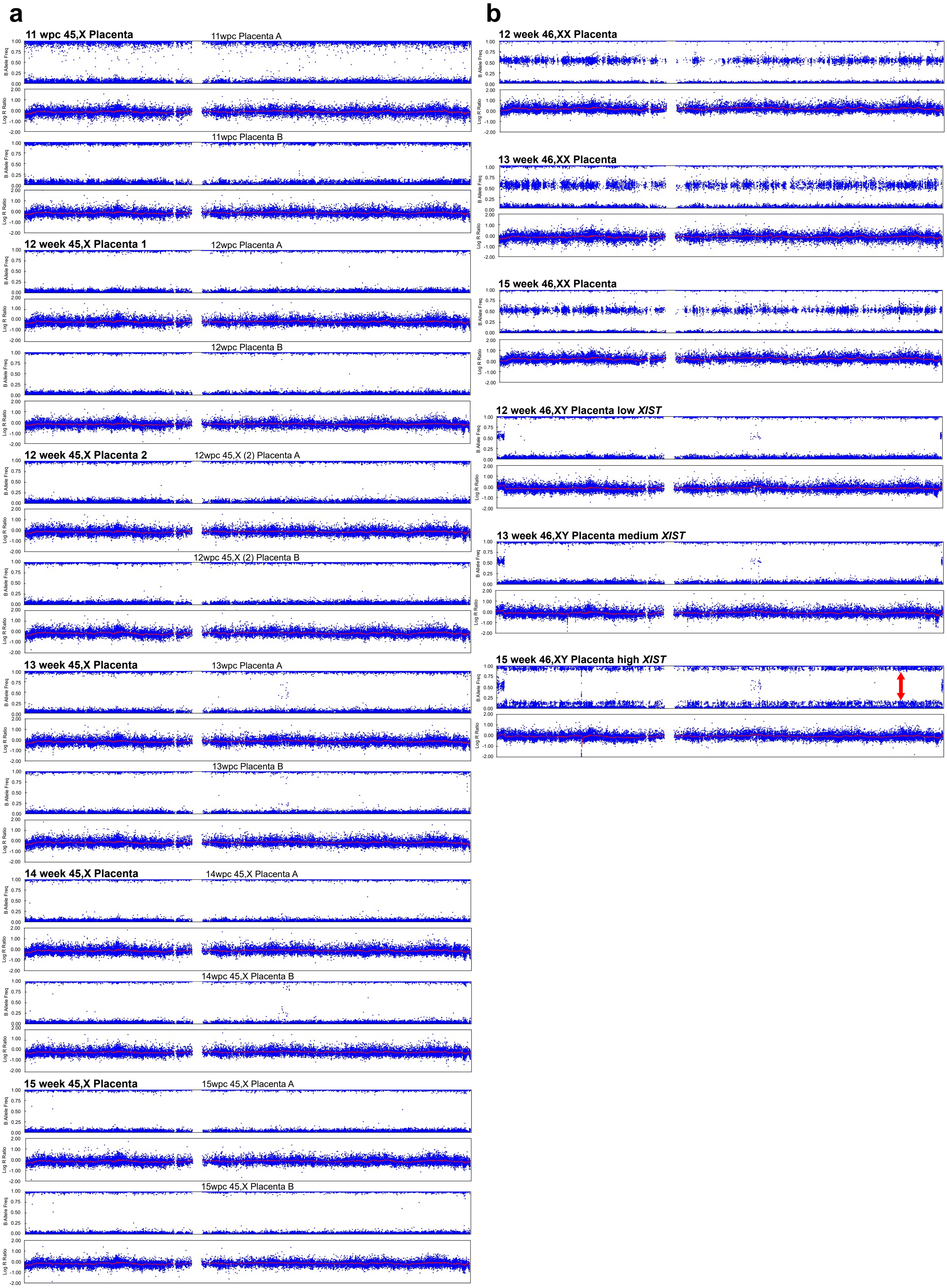

**Supplementary Figure 11. SNP arrays from monosomy X and control placentae for the X chromosome. a** SNP arrays showing the X chromosome in biological replicates taken from each monosomy X placenta (n=12 samples, n=6 placentae). The biological replicates are labelled “Placenta A” and “Placenta B”. **b** SNP arrays showing representative examples of the X chromosome from 46,XX and 46,XY karyotype placentae from 12 weeks post conception (wpc), 13wpc and 15 wpc. For 46,XY placentae, examples with low, medium and high *XIST* counts are shown. For all these SNP profiles, the upper panel shows the B allele frequency, which is the normalized measure of the allelic intensity ratios of two alleles A, B. For B allele frequency, the number of bands seen on the plot minus one usually indicates the number of chromosomes at that given locus. B allele frequencies (BAF) of 0.0, 0.5 and 1.0 are expected in a normal 46,XX sample, representing AA, AB and BB, respectively. The lower panel shows the log R ratio, which is the normalized measure of signal intensity for each SNP marker, as $\log_2$ of the ratio between observed and expected for two copies of the genome. For the log R ratio, a signal clustering around zero shows when the region has two copies; higher or lower signal intensities indicate when there are more or less copies in a genomic region, respectively. The mean value is shown by the smoothened red line indicating the mean value of the chromosome at that position. The presence of a 46,XX cell component in the 46,XY sample with high *XIST* is shown by the arrow.

### Supplementary Figure 12

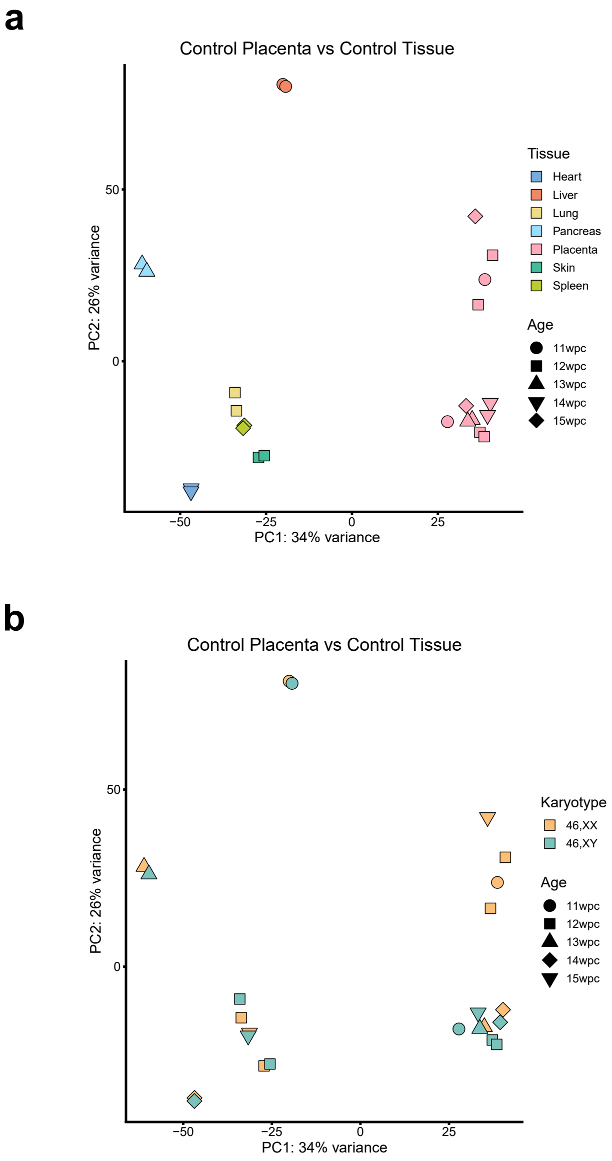


**Supplementary Figure 12. Principal component analysis (PCA) of 46,XX and 46,XY placental samples and control tissue samples. a** PCA showing all 12 placental samples and the different control tissues. **b** PCA showing sample karyotype. PC, principal component; wpc, weeks post conception.

### Supplementary Figure 13

**a**

*AADACL3*

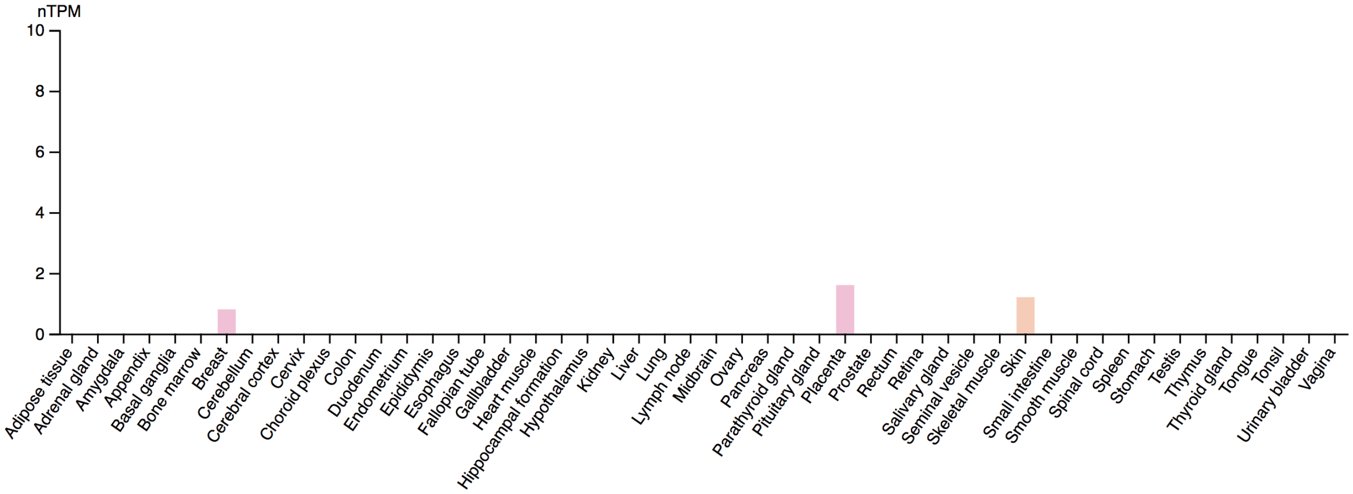


**b**

*CSF2RA*

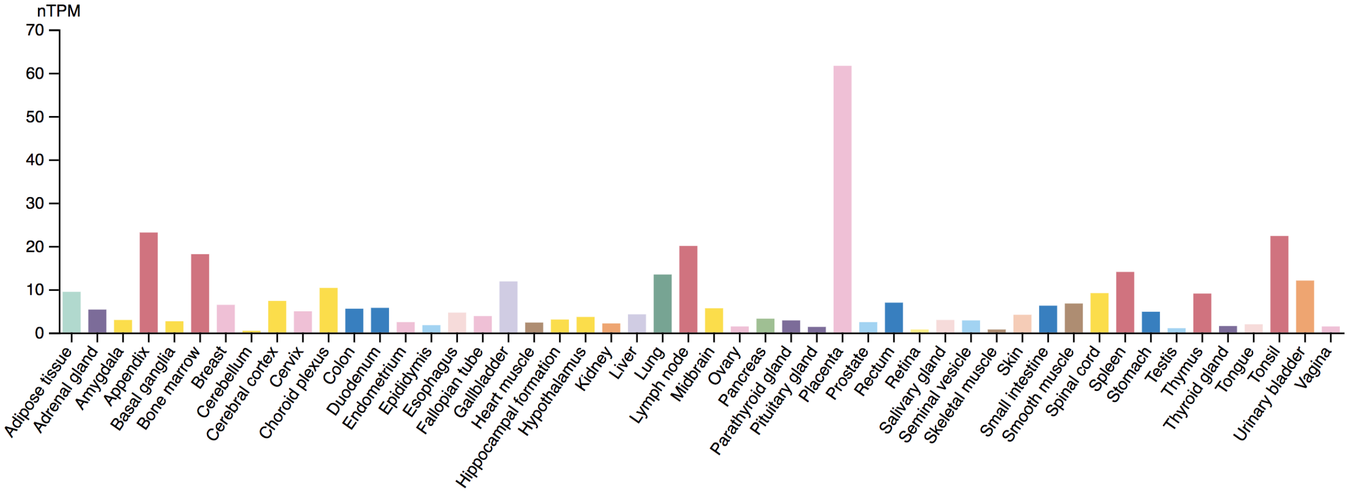


**Supplementary Figure 13. *AADACL3* and *CSF2RA* expression in the Human Protein Atlas Consensus dataset.** Data represent bulk RNA-sequencing expression in a range of adult tissues. Image credit: Human Protein Atlas. **a** *AADACL3* expression (https://www.proteinatlas.org/ENSG00000188984-AADACL3/tissue). **b** *CSF2RA* expression (https://www.proteinatlas.org/ENSG00000198223-CSF2RA/tissue). Data accessed 10/17/23; Protein Atlas version 23.0 available from v23.0.proteinatlas.org (Uhlén et al.,2015)[58] (CC-BY-4.0). nTPM, normalized transcripts per million.

### Supplementary Figure 14

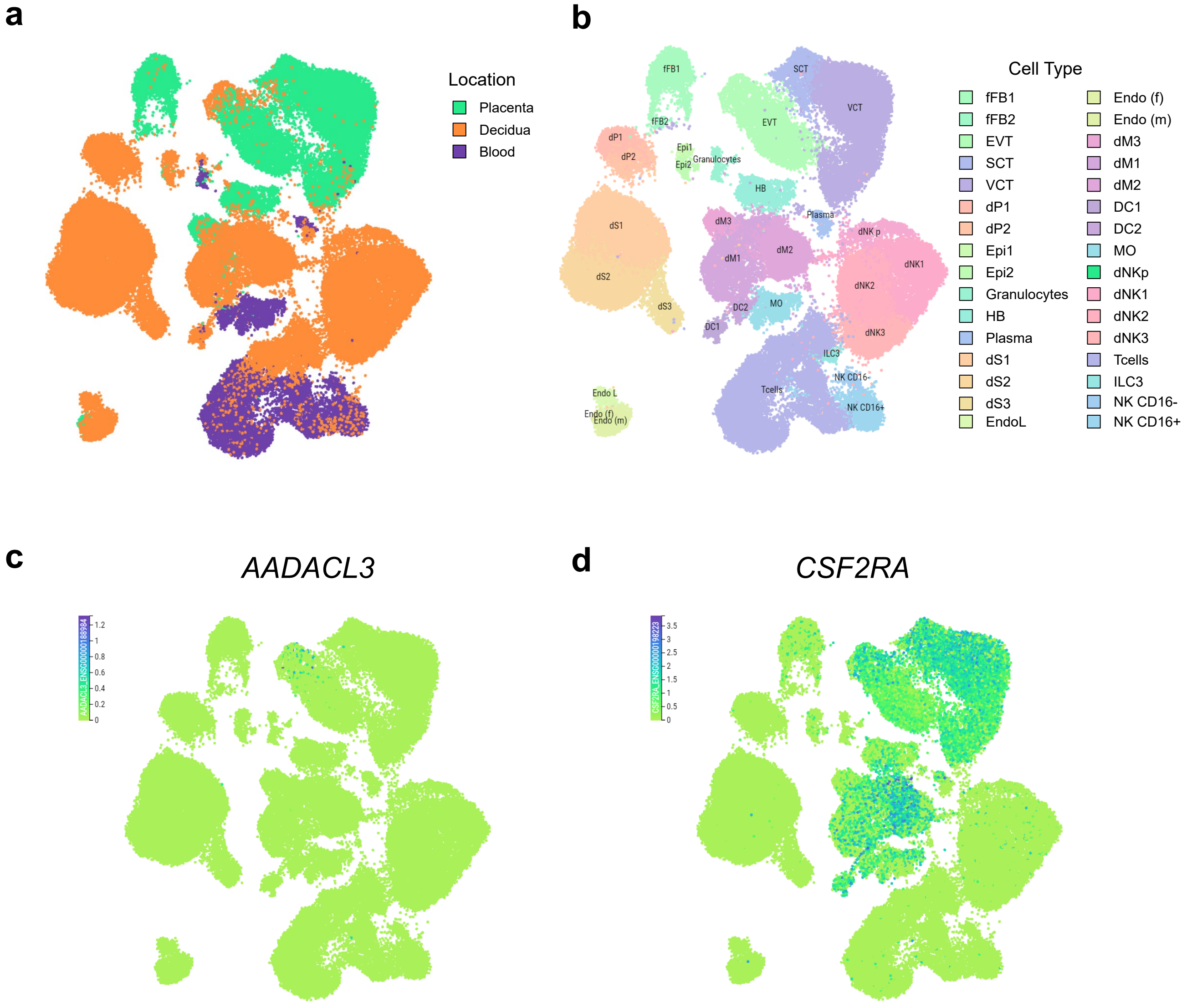


**Supplementary Figure 14. Placental single cell RNA-sequencing expression of *AADACL3* and *CSF2RA* in first trimester placenta. a** Uniform manifold approximation and projection (UMAP) for major tissue clusters. **b** Annotated UMAP key for clusters based on cell type. **c** Feature plot for *AADACL*. **d** Feature plot for CSF2RA. These data were generated by the Vento-Tormo/Teichmann groups at the Wellcome Sanger Institute, Hinxton, UK and can be accessed using CZ CELLxGENE from https://maternal-fetal-interface.cellgeni.sanger.ac.uk/ (Vento-Tormo, R., Efremova, M., Botting, R.A. et al. Single-cell reconstruction of the early maternal–fetal interface in humans. Nature 563, 347–353 (2018); https://doi.org/10.1038/s41586-018-0698-6). This graphic is generated under a Creative Commons Attribution-BY 4.0 International License (https://creativecommons.org/licenses/by/4.0/). DC, dendritic cells; dM, decidual macrophages; dS, decidual stromal cells; Endo, endothelial cells; Epi, epithelial glandular cells; EVT, extravillous trophoblast; f, fetal; F, fibroblasts; HB, Hofbauer cells; ILC, innate lymphocyte cells; l, lymphatic; m, maternal; M3, maternal macrophages; PV, perivascular cells; p, proliferative; SCT, syncytiotrophoblast; VCT, villous cytotrophoblast. CZ CELLxGENE Discover. Retrieved (September 2023), from https://cellxgene.cziscience.com/ (CZ CELLxGENE Discover: A single-cell data platform for scalable exploration, analysis and modelling of aggregated data CZI Single-Cell Biology, et al. bioRxiv 2023.10.30; doi: https://doi.org/10.1101/2023.10.30.563174).

Supplementary Figure 15

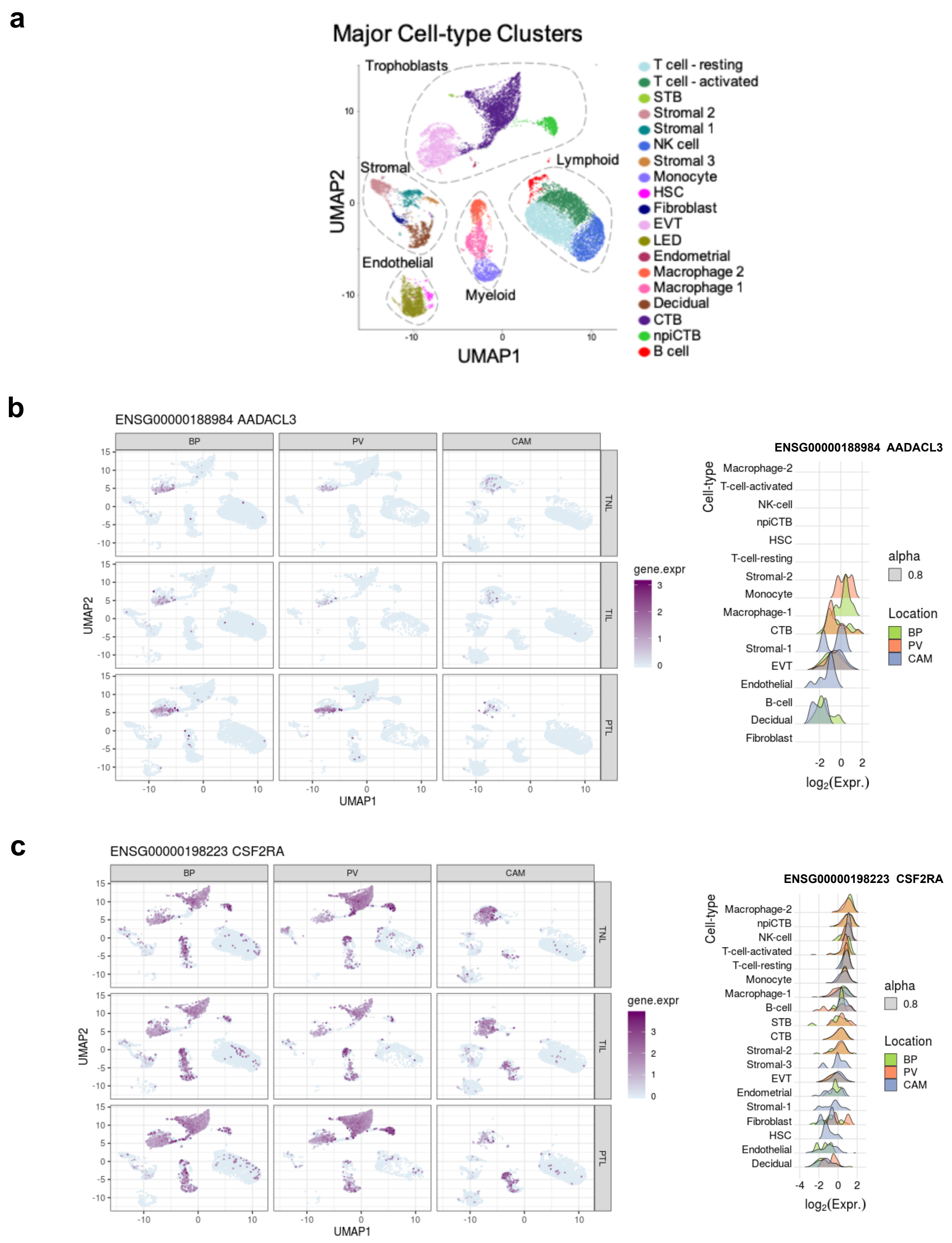


**Supplementary Figure 15. Placental single cell RNA-sequencing expression of *AADACL3* and *CSF2RA* at parturition. a** Annotated uniform manifold approximation and projection (UMAP) for major cell clusters. Reproduced from Pique-Regi et al., 2019 (Pique-Regi, R. et al. Single cell transcriptional signatures of the human placenta in term and preterm parturition. Elife 8, (2019) (CC0 1.0 Universal). **b** UMAP and ridge plots for *AADACL3*. **c** UMAP and ridge plots for *CSF2RA*. Data for panels b and c were accessed from http://placenta.grid.wayne.edu/. BP, basal plate; CAM, chorioamniotic membranes; CTB, cytotrophoblast; EVT, extravillous trophoblast; HSC, hematopoietic stem cell; LED, lymphoid endothelial decidual cell; PTL, preterm labor; PV, placental villi; STB, syncytiotrophoblast; npiCTB, non-proliferative interstitial cytotrophoblast; TIL, term in labor; TNL, term no labor.
